# Drowsiness alters the neural dynamics but not the core computations of multisensory integration

**DOI:** 10.64898/2026.01.12.699055

**Authors:** Clara Alameda, Chiara Avancini, Daniel Sanabria, Tristan A. Bekinschtein, Luis F. Ciria

## Abstract

Fluctuations in alertness shape perception and behaviour, yet how they affect the brain’s ability to integrate information across senses remains poorly understood. Here we investigated whether multisensory integration is preserved as humans transition from wakefulness to sleep. Participants (18 females, 8 males) performed an audio–tactile detection task while electroencephalography tracked spontaneous declines in alertness. Behaviourally, multisensory stimulation continued to facilitate responses during drowsiness, despite slowing and increased omissions. Although race-model predictions were consistent with preserved evidence for multisensory integration, event-related signatures of multisensory interactions were attenuated or no longer detectable during drowsiness. Crucially, multivariate decoding showed that multisensory neural representations remained detectable, although less temporally stable, and early cross-state generalization revealed shared representational structure across alertness levels, indicating preserved core multisensory computations. These findings show that multisensory integration remains functional but dynamically altered as alertness declines, revealing how the brain maintains adaptive behaviour during the wake-to-sleep transition.

**Significance Statement:** Multisensory integration has often been described as a stable and automatic perceptual process. However, growing evidence indicates that it can be modulated by ongoing brain states and cognitive demands. Here, we ask whether this computation remains invariant as alertness declines. We show that multisensory integration is neither abolished nor fully invariant during drowsiness. Instead, behavioural benefits are maintained despite less stable neural dynamics. Although event-related responses are attenuated, core multisensory representations remain shared across levels of alertness. These findings indicate that multisensory integration is preserved across fluctuations in alertness, while their neural dynamics are flexibly altered as alertness decreases.

## INTRODUCTION

During a night shift, a doctor hears a beep and feels her pager vibrate on her belt, signals that would usually prompt her to confirm the call and head to the emergency room. Whether and how quickly that response unfolds likely depends on how awake she is at that moment. In daily life, we experience natural downward shifts in alertness that measurably affect perception, decision making and action (Lacaux et al., 2024). Drowsiness can be understood as a transitional state between attentive wakefulness and sleep that often preserves behavioural responsiveness, providing a useful paradigm to identify which cognitive operations persist and which begin to fragment as alertness wanes (Goupil & Bekinschtein, 2012).

Neuroimaging research has begun to characterize how transitions into drowsiness affect cognition (Goupil & Bekinschtein, 2012; Lacaux et al., 2024). Early sensory processing appears relatively preserved: auditory evoked responses such as the N1 remain detectable, though often delayed (Strauss et al., 2015). Simple target detection also persists, albeit with slower responses and occasional lapses (Noreika et al., 2020; Xu et al., 2023). As task demands increase, performance deteriorates, with rising error rates and volatility (Ciria et al., 2021; Xu et al., 2023). Not all functions decline uniformly, however, and preserved performance has been linked to compensatory or reconfigured neural dynamics that sustain behaviour under reduced alertness (Canales-Johnson et al., 2020; Jagannathan et al., 2022).

Although progress has been made in characterizing sensory and cognitive processing during the wake–sleep transition, most research has relied on unisensory paradigms, which do not capture the multisensory nature of everyday behaviour. Even simple responses depend on whether events occur in one modality or across several. When redundant multisensory signals are presented, reaction times are typically faster than to either unisensory signal alone, a robust phenomenon known as the redundant-signals effect (RSE; Meredith & Stein, 1983).

In humans, multisensory integration is often operationalized using an additive model, in which the neural response to a multisensory stimulus is compared with the sum of its unisensory components; deviations from additivity are taken as evidence of cross-modal interaction (Besle et al., 2004). Studies in alert participants report heterogeneous neural signatures, including early superadditivity, later subadditivity, or mixed effects across time windows and regions (e.g., Quinn et al., 2014; Senkowski et al., 2011). Despite extensive work under wakeful conditions, little is known about how these behavioural and neural markers are influenced by fluctuating alertness.

Could drowsiness alter how the doctor integrates the pager’s vibration and beep, thereby slowing or degrading her response? Evidence from non-wakeful states is limited and mostly indirect. Integration effects have been observed in multisensory neurons under anaesthesia in animal models (Stein & Stanford, 2008), whereas pharmacologically induced unconsciousness can diminish bimodal cortical responses (Ishizawa et al., 2016). The only study examining multisensory detection at sleep onset reported preserved audiovisual reaction times during drowsiness, suggesting redundancy-based compensation for reduced alertness detriment in performance, but did not assess computational or neural markers of integration (Artigas et al., 2025). Thus, a comprehensive characterization of multisensory integration and its neural readouts during natural drifts into drowsiness is still missing.

In this pre-registered study (https://osf.io/mu6ne), we addressed this critical gap by asking healthy adults to perform an audio–tactile detection task while EEG tracked spontaneous transitions between wakefulness and light sleep. We combined behavioural measures, race-model analyses, event-related potentials and multivariate decoding to test whether multisensory redundancy benefits persist as alertness declines and whether they rely on comparable neural mechanisms across states. Based on prior evidence for robust multisensory enhancement in wakefulness and relative preservation of early sensory processing during drowsiness, we predicted that behavioural redundancy benefits would remain detectable during drowsiness, whereas the neural dynamics supporting multisensory integration would become delayed and less stable, revealing how core multisensory computations are preserved yet altered as alertness wanes.

## MATERIALS AND METHODS

### Participants

We recruited 30 young adults through local advertisements and the research centre’s participant platform. Twenty-six participants (18 females; 18–32 years) were included in the final analyses. Three were excluded due to EEG acquisition problems and one due to prolonged non-responsiveness (hit rates <50% in some conditions). To facilitate sleep onset during the experimental session, we recruited healthy individuals with Epworth Sleepiness Scale scores between 7 and 14 (Johns, 1991). This range reflects a tendency toward increased sleep propensity, typically corresponding to mild levels of daytime sleepiness that do not necessarily indicate the presence of a sleep disorder (Johns, 1992; Ohayon, 2008). All participants reported normal hearing and no history of sleep, neurological or psychiatric disorders. Participants received monetary compensation. The study was approved by the Ethics Committee of the University of Granada (3508/CEIH/2023).

### Experimental task

Participants performed an audio–tactile detection task adapted from Murray et al. (2005). Stimuli were auditory, tactile, or spatially aligned audio–tactile pairs presented on the left or right, yielding six configurations (auditory left/right; tactile left/right; audio–tactile left/right). Trials were randomized with equal probability (330 per configuration; 1980 total). The inter-stimulus interval varied between 2–4 s and the response window was 4 s. No feedback was provided. Participants responded as quickly as possible to any stimulus by pressing a button with the thumb of their dominant hand. Before the main task, participants completed a 24-trial familiarization block (6 trials per configuration) to confirm detectability and spatial discriminability.

Auditory stimuli were 30-ms white-noise bursts (∼60 dB) delivered via two loudspeakers positioned in front of the participant and separated by 100° in azimuth (Amazon Basics, Amazon.com, Inc., Seattle, WA, USA). Tactile stimuli were 30-ms vibrations (∼100 Hz) delivered to the left or right upper arm (midpoint between shoulder and elbow) using miniature stimulators (“Tactor”; Dancer Design, St Helens, United Kingdom; 18-mm diameter). Participants wore earplugs to ensure tactile stimulation was inaudible while maintaining audibility of the acoustic stimuli (Moldex-Metric, Culver City, CA, USA). This was explicitly addressed in the task instructions and verified during a familiarization phase prior to the experiment, during which correct earplug placement and the inaudibility of the tactile stimulation were confirmed. The familiarization was repeated if necessary. In addition, between blocks, we ensured that the earplugs remained properly fitted and comfortable. Stimulus presentation and response collection were implemented in MATLAB (MathWorks, Natick, MA, USA) using Psychtoolbox (Kleiner et al., 2007) and delivered via a custom-built audio-tactile response box (CRAT) using hardware-level relays and Arduino-based control. Temporal precision was verified using an oscilloscope, confirming millisecond-level synchronization across auditory, tactile and trigger signals. The loudspeakers were positioned approximately 50 cm from the participant’s head, and the small delay introduced by sound propagation (∼1.5 ms) was estimated and compensated to ensure temporal alignment between auditory and tactile stimuli at the point of perception.

### Procedure

Participants completed a single experimental session between 2 pm and 5 pm after a normal night’s sleep. Following informed consent and task instructions, participants were fitted with a 64-channel EEG cap (EASYCAP GmbH, Herrsching, Germany). The experiment was conducted in a darkened room, with participants seated in a reclined chair, with their eyes closed, and were instructed to minimize movement while responding as quickly as possible.

The task consisted of alternating alert and drowsy blocks. In alert blocks (240 trials, ∼15 min), participants were instructed to remain awake and attentive. In drowsy blocks (750 trials, ∼45 min), they were allowed to fall asleep. If they became unresponsive, the experimenter gently woke them by knocking at the door. To facilitate the transition to sleep, participants were made comfortable (e.g., reclining position, blanket, removal of shoes). This alert–drowsy sequence was repeated twice to maximize the number of trials obtained under reduced alertness, following established protocols (Canales-Johnson et al., 2020; Ciria et al., 2021; Jagannathan et al., 2022). Between blocks, short breaks (∼5 min) were introduced to promote re-alerting. During these intervals, lights were turned on, the chair was brought to an upright position, and participants briefly interacted with the experimenter and completed short subjective reports of their ongoing experience. These procedures are known to transiently restore alertness at the beginning of the block (e.g., Helton & Russell, 2015; Ross et al., 2014; Schumann et al., 2022). The full session lasted approximately 3 hours (1980 trials in total).

### Alertness classification

Given that the wake-to-sleep transition is characterised by a progressive attenuation of the EEG alpha rhythm together with an increase in theta activity (De Gennaro et al., 2001), drowsiness was quantified at the single-trial level using spectral power estimates, as implemented in previous studies (Bareham et al., 2014; Ciria et al., 2021; Comsa et al., 2019; Xu et al., 2023). For each trial, spectral power was computed in the prestimulus interval (−2000 to 0 ms relative to target onset) using a continuous wavelet transform (3 cycles at 3 Hz to 8 cycles at 40 Hz). Theta (4–6 Hz) and alpha (10–12 Hz) power were averaged across central (C3, C2) and occipital (O1, Oz, O2) electrodes, respectively. A theta/alpha ratio was then computed for each trial, yielding a single index of alertness.

Trials were initially classified for each participant using percentile-based thresholds, with the highest 33% labelled as drowsy and the lowest 33% as awake. This approach captures a continuous process by defining two relatively meta-stable alertness states (regimes) and is commonly used in studies of the wake–sleep transition (e.g., Bareham et al., 2014, 2015; Ciria et al., 2021; Xu et al., 2025). To reduce spurious fluctuations and account for temporal continuity, a smoothing step based on sleep hysteresis principles was applied (Saper et al., 2010), whereby isolated trials were relabelled to match the surrounding state when embedded within sustained periods (≥10 trials). In addition, the first 100 trials of each block were conservatively labelled as awake, in line with previous studies (e.g., Ciria et al., 2021), to account for the re-alerting effect induced by breaks (see Procedure). Importantly, final labels were determined exclusively by the EEG-based theta/alpha ratio and were independent of block structure (see Fig. 1C). Behavioural measures (reaction times, omissions) were not used for classification but analysed independently. Blocks were designed to promote repeated transitions between alert and drowsy periods within participants. The validity of this approach is supported by its correspondence with behavioural markers and by additional control analyses reported in the Supplemental Material (Text S1). Specifically, we assessed the robustness of the theta/alpha-based classification by recomputing the index after separating the aperiodic component and individualizing the alpha band based on each participant’s peak frequency (as suggested during peer-review), as well as by inspecting the temporal evolution of the ratio alongside behavioural measures at the single-participant level (see Supplemental Fig. S1-S2; complete participant-level plots available in the study OSF repository https://osf.io/7pt86/files/osfstorage).

**Figure 1.**
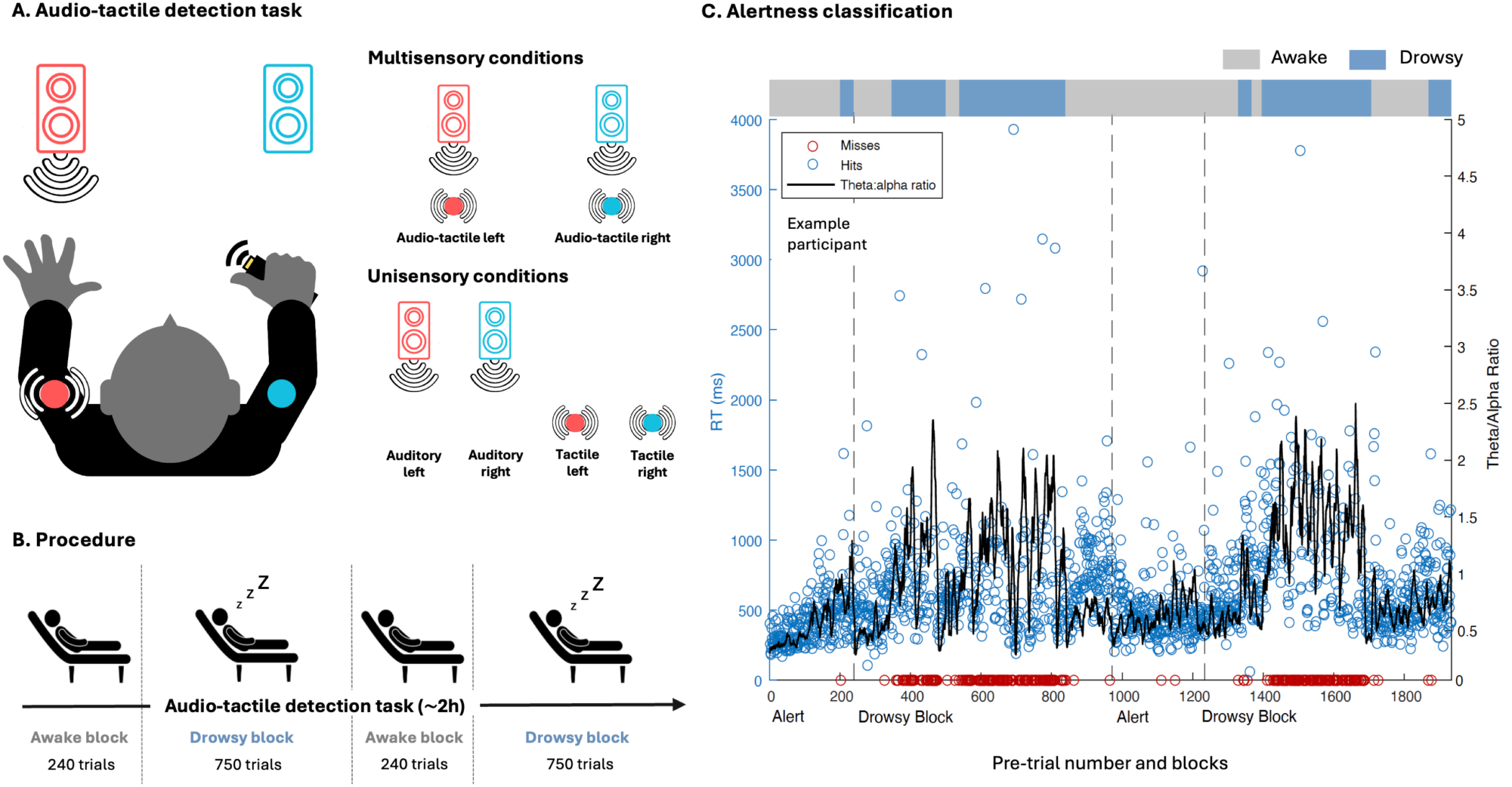
Experimental design and theta/alpha ratio-based alertness classification. A) Schematic representation of the audio-tactile detection task implemented. Participants were asked to respond as fast as possible with the dominant hand to randomized stimulus conditions: unisensory stimulus, tactile (vibration on the arm) or auditory (white noise bursts through loudspeakers), and multisensory stimuli (auditory and tactile simultaneous and spatially aligned stimuli). B) Schematic representation of experimental procedure. Participants performed the experimental task, divided into four blocks. In the short “awake” blocks, participants were asked to stay alert and focused on the task, while in the longer “drowsy” blocks, participants were explicitly told that they could fall asleep, yet if they stopped responding for five consecutive trials or more, they would be awakened by the experimenter. These two types of blocks were alternated during the session. C) Automatic classification of alertness level during the experimental session (representative participant). The black line depicts changes in the theta/alpha ratio (central-occipital electrodes clusters) during the pre-trial period (2 s before the stimulus onset). The horizontal bars on top represent trials classified as awake (grey) or drowsy (blue). The variability in the reaction times (blue circles) closely follows the changes in theta/alpha ratio. Notice that circles on the horizontal axis (reaction time equal to zero) were non-responsive trials, usually during reduced alertness periods.

### EEG acquisition and preprocessing

EEG data were collected using a 64-channel actiCHamp amplifier with Brain Vision software. Data were sampled at 1000 Hz and referenced to FCz according to the extended 10–20 system. Electrode impedances were kept below 10 kΩ. Continuous data were downsampled to 250 Hz, band-pass filtered between 0.5 and 40 Hz using a zero-phase finite impulse response (FIR) and line noise was attenuated using a 50 Hz FIR band-stop filter. Peri-auricular electrodes (FT9, FT10, TP9, TP10) were excluded and ocular artifacts were removed using Independent Component Analysis (ICA). Data were epoched from −200 to 1500 ms relative to stimulus onset. Trials rejected in behavioural analyses and those exceeding ±150 μV were removed. Signal was re-referenced to the average. After preprocessing, an average of 789.69 (SD = 67.64) trials per participant were retained in the awake condition, and 562.62 (SD = 51.7) in the drowsy condition. Preprocessing was conducted with EEGLAB (Delorme & Makeig, 2004) and FieldTrip (Oostenveld et al., 2011).

### Experimental Design and Statistical Analysis

The study employed a within-subject design with two levels of alertness (awake, drowsy) and three stimulus types (auditory, tactile, audio–tactile). All participants completed all conditions. Alertness was classified at the single-trial level as described above.

#### Behavioural reaction times and response rate analyses

To investigate whether the magnitude of multisensory temporal benefits (i.e., RSE [redundant signals effect]) is modulated during the wake-sleep transition, mean reaction times (RTs) and response rates (i.e., 0 meaning no response and 1 meaning the participant responded to every trial) were computed for each stimulus type and alertness state. Missed trials and RTs below 150 ms were excluded. RTs were analyzed using linear mixed-effects models (*lme4* R package; Bates et al., 2014) with participant as a random factor and alertness (awake, drowsy), type of stimuli (tactile, auditory, audiotactile), side (left, right) and their interaction as fixed effects. Model selection was based on AIC, starting from the maximal model and removing predictors that did not improve fit (see Text S1 for details on AIC values). Significant effects were followed by Bonferroni-corrected post hoc comparisons derived from the fitted linear mixed-effects models using estimated marginal means (*emmeans*). Degrees of freedom were estimated using Satterthwaite’s approximation, as implemented in the lmerTest package (Kuznetsova et al., 2017). Equivalent models were fitted to response rate data.

As suggested during the peer review process, we repeated the main analyses using log-transformed RTs, to assess robustness. This transformation improved residual normality and homoscedasticity without altering the pattern of results. Additionally, we examined whether awake trials differed depending on the block context in which they occurred. Trials were classified into three categories: awake trials within awake blocks, awake trials within drowsy blocks, and drowsy trials. Behavioural measures (RT and response rate) were analysed using linear mixed-effects models analogous to the main analyses. Full details and results are reported in the Supplemental Material (see Text S1, “Control analyses for alertness classification based on the theta/alpha ratio”).

#### Computational race modelling

Each participant RT distributions were further analysed to evaluate different hypotheses accounting for potential multisensory interactions. To do so, we implemented Innes & Otto’s (2019) approach via the modelling package Matlab RSE-box. After excluding misses and anticipations, we transformed the RTs into rates (1/RT) and excluded data points that deviated by more than 3 SDs. Cumulative distribution functions (CDFs) were computed using 50 quantiles (i.e., time bins) per participant, condition, and alertness state.

First, empirical redundancy benefits were estimated as the area between the multisensory CDF and the faster unisensory CDFs (Grice’s bound; see Figure 2C). Empirical RSE benefits were compared between alertness states using two-tailed paired-samples t-tests, as paired differences did not deviate significantly from normality (Shapiro–Wilk test, *p* = 0.11).

**Figure 2.**
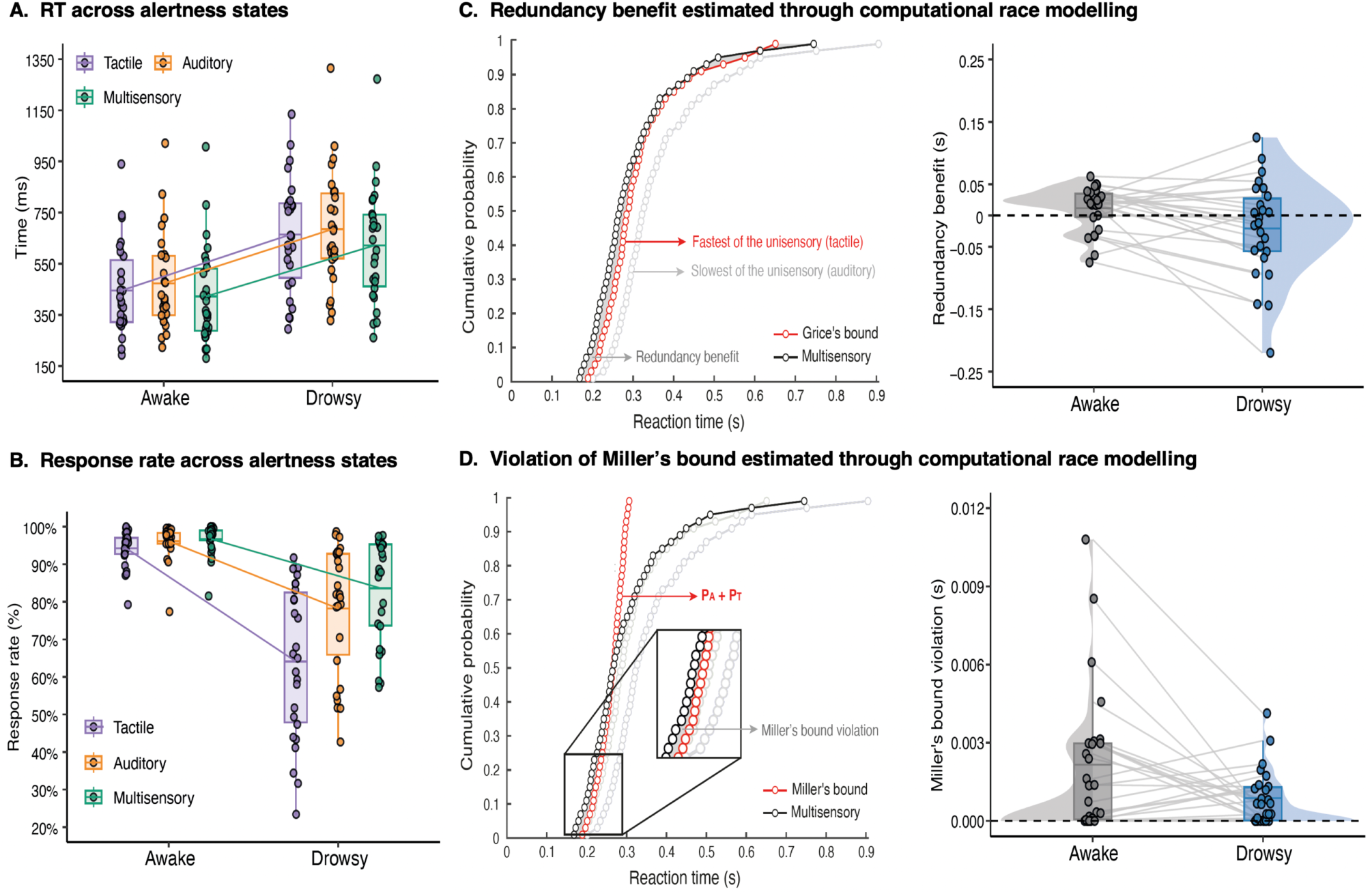
The impact of drowsiness on redundancy effect indices. A) RTs across stimulus conditions under the two alertness states. Each dot represents a participant’s mean RT for that condition. The central line of the boxplot depicts the mean, the box spans the twenty-fifth to the seventy-fifth percentile, and upper and lower whiskers indicate a distance of 1.5 times the interquartile range above the percentiles Coloured lines link the mean values between Awake and Drowsy for each stimulus, illustrating the change across conditions. Although responses were consistently slower during drowsiness across all stimulus conditions, linear mixed-effects models revealed that the classical RSE was preserved during drowsiness, with shorter RTs for multisensory versus each unisensory condition. B) Response rate across stimulus conditions and alertness states. While response rates were near ceiling during wakefulness across all stimulus conditions, they dropped markedly during drowsiness, particularly for the tactile condition. C) *Left:* empirical redundancy gain of an example participant, quantified as the area between the multisensory CDF and Grice’s bound (i.e., the fastest of the unisensory RT distribution). Each dot represents one of the 50 time bins into which the participant’s RT were divided for each stimulus condition. *Right:* Group-level comparison of the redundancy benefit between alertness states. Grey lines connect the two mean values (dots) from the same subject across alertness states. No differences were found in the empirical benefit between alertness states. D) *Left:* Miller’s bound violation of an example participant, quantified as the area where the multisensory CDF exceeds Miller’s bound (i.e., the maximum predicted by statistical facilitation), revealing multisensory integration. Miller’s violation was present in both alertness states. However, the difference between wakefulness and drowsiness did not reach statistical significance.

Second, to test whether multisensory speed-ups could be explained by statistical facilitation alone (i.e., responses triggered by whichever unisensory channel finishes first, assuming independent channels), we evaluated Miller’s race model inequality (1982):

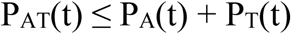

With P being the cumulative probability of responding by time *t* for an audio-tactile signal (AT), an auditory signal (A) or a tactile signal (T). Violations of Miller’s bound were quantified as the area where the multisensory CDF exceeded the summed unisensory CDFs (see Figure 2D). Given that violation values are bounded at zero and deviate from normality, these were tested using non-parametric Wilcoxon signed-rank tests, both against zero and between alertness states. To control for multiple comparisons across the two primary indices (Grice’s bound and Miller’s bound violation), a Bonferroni-corrected significance threshold of α = 0.025 was applied. Correlation analyses were additionally performed to assess the relationship between individual Miller’s bound violation values across wakefulness and drowsiness using Spearman’s rank correlation.

In addition, to address potential alternative explanations of race model violations, we conducted control analyses examining the influence of trial-history effects, global RT slowing, and changes in unisensory processing. Trial-history effects (e.g., modality or response side switching between consecutive trials) can affect reaction times and bias estimates of multisensory gain (Shaw et al., 2020), while changes in unisensory response speed may alter RT distributions and influence the likelihood of observing race model violations, particularly under conditions of overall slowing (Couth et al., 2018). We therefore quantified switching costs and tested whether these, together with global slowing and unisensory RT changes, accounted for variance in violation across participants (see Supplemental Fig. S4).

#### ERP analyses

Trials were grouped by alertness (awake, drowsy) and type of stimuli (auditory, tactile and audio-tactile), resulting in 6 conditions: awake tactile, awake auditory, awake multisensory, drowsy tactile, drowsy auditory and drowsy multisensory. Trial counts were equalized across stimulus types within each state by random subsampling. Following this procedure, an average of 235.85 (SD = 27.88) trials per participant and stimulus type were computed in the awake condition, and 117.04 (SD = 36.09) in the drowsy condition. ERPs were computed from 0–500 ms using a −200 ms baseline.

Multisensory effects were assessed using the additive model (Berman, 1961; Besle et al., 2004), which assumes that if auditory and tactile signals are processed independently, the ERP elicited by their combined presentation should equal the arithmetic sum of the corresponding unisensory ERPs. Deviations from this summed response (i.e., superadditivity or subadditivity) are therefore taken as evidence of cross-modal neural interactions. For each alertness state, auditory and tactile ERPs were summed and compared against multisensory ERPs. Cluster-based permutation tests (Maris & Oostenveld, 2007) identified significant spatiotemporal differences without predefined temporal or spatial windows (*p* < 0.05; 5000 permutations). Direct differences between alertness states were evaluated using difference-of-differences contrasts within the same permutation framework.

As an additional analysis suggested during peer review, peak latencies were extracted from the multisensory interaction waveform (multisensory-sum of unisensory responses), focusing on components associated with multisensory integration effects (i.e., non-additive response magnitudes) and compared using paired-sample *t*-tests (see Text S1).

#### Multivariate pattern analysis (MVPA) analyses

We used Linear Discriminant Analysis (LDA) to decode whether activity reflected the multisensory condition versus the arithmetic sum of the unisensory conditions, in a stimulus-locked window (−200 to 500 ms) with MVPAlab (López-García et al., 2022). Trial numbers were matched across side and class within each fold. To prevent information leakage, z-scaling (mean/SD per electrode) was fitted on training data and applied to both training and test sets. Decoding was computed every three time points, and performance was quantified as Area Under the Curve (AUC).

We used a non-parametric cluster-based permutation test comparing observed decoding curves to an empirical chance distribution built by shuffling trial labels within participants (1000 permutations). Group-level null decoding curves were generated from these permutations, and the 95th percentile was used to set the threshold for significant effects. Cluster size and extent were compared to those from permuted data, and multiple comparisons were controlled using False Discovery Rate (López-García et al., 2022). To probe temporal dynamics, we applied temporal generalization (training at each time point, testing across all) to obtain Temporal Generalization Matrices (TGMs) and evaluated them with the same permutation framework. Cross-state generalization was tested by training in wakefulness and testing in drowsiness (and vice versa)

## RESULTS

To assess the impact of declining alertness on multisensory integration, we combined behavioural, computational, and neural analyses. Alertness state was defined at the single-trial level using established prestimulus EEG markers of the wake-to-sleep transition, based on the relative power of theta and alpha rhythms (see Figure 1C). We first examined mean reaction times and response rates to test whether multisensory redundancy benefits are preserved during drowsiness. Because mean reaction times alone cannot dissociate facilitation from integration, we next used race-model analyses to probe changes in multisensory computational efficiency, and EEG-based univariate and multivariate analyses to characterize how the neural dynamics supporting multisensory integration are altered as alertness wanes.

### Multisensory redundancy benefits are preserved during drowsiness despite slower and less reliable responses

We first tested whether alertness modulated the classical redundant signals effect (RSE; i.e., faster responses for the multisensory stimuli vs. the unisensory ones). The average RTs and response rate per participant were calculated for each condition and then fitted using hierarchical linear mixed-effects models. RTs yielded a robust effect of alertness, *F*(1,281) = 416.98, *p* < 0.001, *η²p* = 0.6, and type of stimuli, *F* (2,281) = 10.37, *p* < 0.001, *η²p* = 0.07. Bonferroni-corrected post hoc contrasts derived from the fitted model (estimated marginal means) showed significantly faster responses for multisensory than auditory stimuli, *t*(281) = 4.54, *p* < 0.001, *d* = -0.62, and vs. tactile stimuli, *t*(281) =-2.61, *p* = 0.02, *d* = -0.36, but no differences between the two unisensory conditions (auditory vs. tactile), *t*(281) =1.92, *p* =0.16, *d* = 0.27 (see Figure 2A). Importantly, no reliable interaction between alertness state and stimuli was observed *F*(2,281) = 0.29, *p* = 0.766, *η²p* = 0.002. Most participants showed a RSE benefit in mean RT, with 19 out of 26 participants exhibiting faster multisensory responses during wakefulness and 16 during drowsiness. Analyses on log-transformed RTs yielded the exact same pattern of main effects and post hoc contrasts (see Text S1).

For response rate, linear mixed-effects models showed a main effect of alertness, *F*(1, 286) = 325.33, *p* < 0.001, *η²p* = 0.53, and type of stimuli, *F*(2, 286) = 33.81, *p* < 0.001, *η²p* = 0.19, as well as a significant interaction between alertness and stimulus type, *F*(2, 286) = 20.09, *p* < 0.001, *η²p* = 0.12. Bonferroni-corrected post hoc contrasts from the mixed-effects model revealed increased omissions for tactile stimuli compared with auditory stimuli during drowsiness, *t*(297) = -7.07, *p* < 0.001, *d* = -1.41, and compared with multisensory stimuli, *t*(297) = -9.78, *p* < 0.001, *d* = -1.96, but not during wakefulness (awake tactile vs. awake auditory: *t*(297) = -0.95*, p* = 1.000, *d* = -0.19; awake tactile vs. awake multisensory: *t(*297) = -1.25, *p* = 1.000, *d* = -0.25; see Figure 2B).

Additional analyses comparing awake trials across block contexts revealed a graded effect on RTs, with slower responses for awake trials embedded within drowsy blocks. Crucially, however, RTs in both awake conditions remained significantly faster than in drowsy trials, indicating higher levels of alertness despite differences in time-on-task. As expected, response rates did not differ between awake conditions but were markedly reduced in drowsy trials (see Supplemental Fig. S3). Full statistical results are reported in the Supplemental Material (Text S1).

### Computational evidence for multisensory integration is preserved during drowsiness

In the next analyses, we focused on computational *race models*, extensively used in the multisensory integration literature (e.g., Innes & Otto, 2019; Mercier & Cappe, 2020; Murray et al., 2005). They allow for subject-by-subject assessment of whether the RT distribution in the multisensory condition can be explained by an interaction between the two unisensory signals when presented simultaneously, yielding a fine-grained assessment of redundancy beyond mean RT speedups. First, we examined the empirical RSE benefit, defined as the RT acceleration observed when stimuli are presented across multiple sensory modalities compared to a single modality. To quantify this benefit, we calculated the area between the cumulative distribution function (CDF) for the multisensory condition and the fastest of the two unisensory CDFs (the so-called, Grice’s bound). The resulting empirical benefit area was computed for each participant under both alertness states (wakefulness and drowsiness) and compared using two-tailed paired-samples *t*-tests. By examining the entire RT distribution rather than relying solely on mean RT, this analysis provides a more sensitive assessment of the redundancy gain. The analysis did not reveal significant differences in the empirical RSE benefit between alertness states, *t*(25) = -2.36, *p* = 0.026, *d* = 0.54 (Bonferroni-corrected α = 0.025; see Figure 2C), with 19 of 26 participants exhibiting a multisensory benefit in wakefulness and 15 in drowsiness.

Then, to determine whether the observed RSE benefit could be attributed solely to statistical facilitation (i.e., the fastest of the two unisensory signals triggers the response in the multisensory condition), we evaluated the race model inequality (Miller, 1982). According to this approach, the cumulative RT distribution for the multisensory condition should not exceed the sum of the cumulative probability of the unisensory distributions (a sum known as Miller’s bound), assuming no interaction occurs between the sensory signals when they are presented simultaneously. The area of violation of the race model inequality (Miller’s bound violation), accounting for multisensory integration when its value is higher than 0, was estimated per participant and condition. Given that violation values are bounded at zero and deviate from normality, non-parametric Wilcoxon signed-rank tests were used. One-sample tests against zero revealed a robust race model violation in both the awake condition (*V* = 210, *p* < 0 .001, *r* = 0.77) and the drowsy condition (*V* = 190, *p* < 0.001, *r* = 0.75), indicating reliable multisensory integration in both states. In contrast, the comparison between wakefulness and drowsiness did not reach significance (*V* = 212, *p* = 0.079, *r* = 0.32; see Figure 2D), with 20 of 26 participants showing violation values greater than zero during wakefulness and 19 during drowsiness. No significant correlation was observed between Miller’s bound violation values in wakefulness and drowsiness (Spearman: *ρ* = 0.08, *p* = 0.69). Notably, control analyses indicate that these effects are unlikely to be driven by trial-history dependencies, global RT slowing, or changes in unisensory processing (see Text S1).

### ERP signatures of multisensory integration are attenuated during drowsiness

In line with the extensive literature on the neural mechanisms underlying multisensory integration (Stevenson et al., 2014), the ERPs analyses were based on the additive model (Berman, 1961). The evoked potentials elicited by the multisensory stimulus were compared against the algebraic sum of the evoked responses to each unisensory stimulus presented separately, under the premise that differences in the brain responses to simultaneous versus separate signals may indicate the presence of multisensory interactions/integration (see Methods section for further discussion). Although cluster-based permutation tests were conducted across the full 0–500 ms time window without predefined temporal or spatial constraints, we describe the results in terms of early and late intervals for interpretative clarity, in relation to well-established stages of sensory processing and decision-making reported in the literature (e.g., Mercier & Cappe, 2020).

At early processing stages (<150 ms after stimulus onset), cluster-based permutation analyses revealed the classical multisensory superadditivity effect (i.e., an early neural marker of multisensory integration characterized by higher voltage amplitude in the multisensory condition vs. the sum of the unisensory conditions) in frontal-central (44-128 ms, *p* = 0.014; increased negativity for multisensory vs. unisensory) and parietal-occipital electrodes during wakefulness (40-108 ms, *p* = 0.026; increased positivity; see Figure 3A-B, up). Strikingly, this so-called superadditivity ERP marker was not observed in drowsiness (frontal cluster: 112-124 ms, *p* = 0.39; parietal cluster: 108-120 ms, *p* = 0.45; see Figure 3A-B, down). Nevertheless, direct comparison between states (awake vs. drowsy) multi-sum differences did not reach significance before 150 ms, suggesting that while the effect appears to be state-dependent during the encoding stage, statistical evidence for a difference between awake and drowsy conditions at this early stage is lacking.

**Figure 3.**
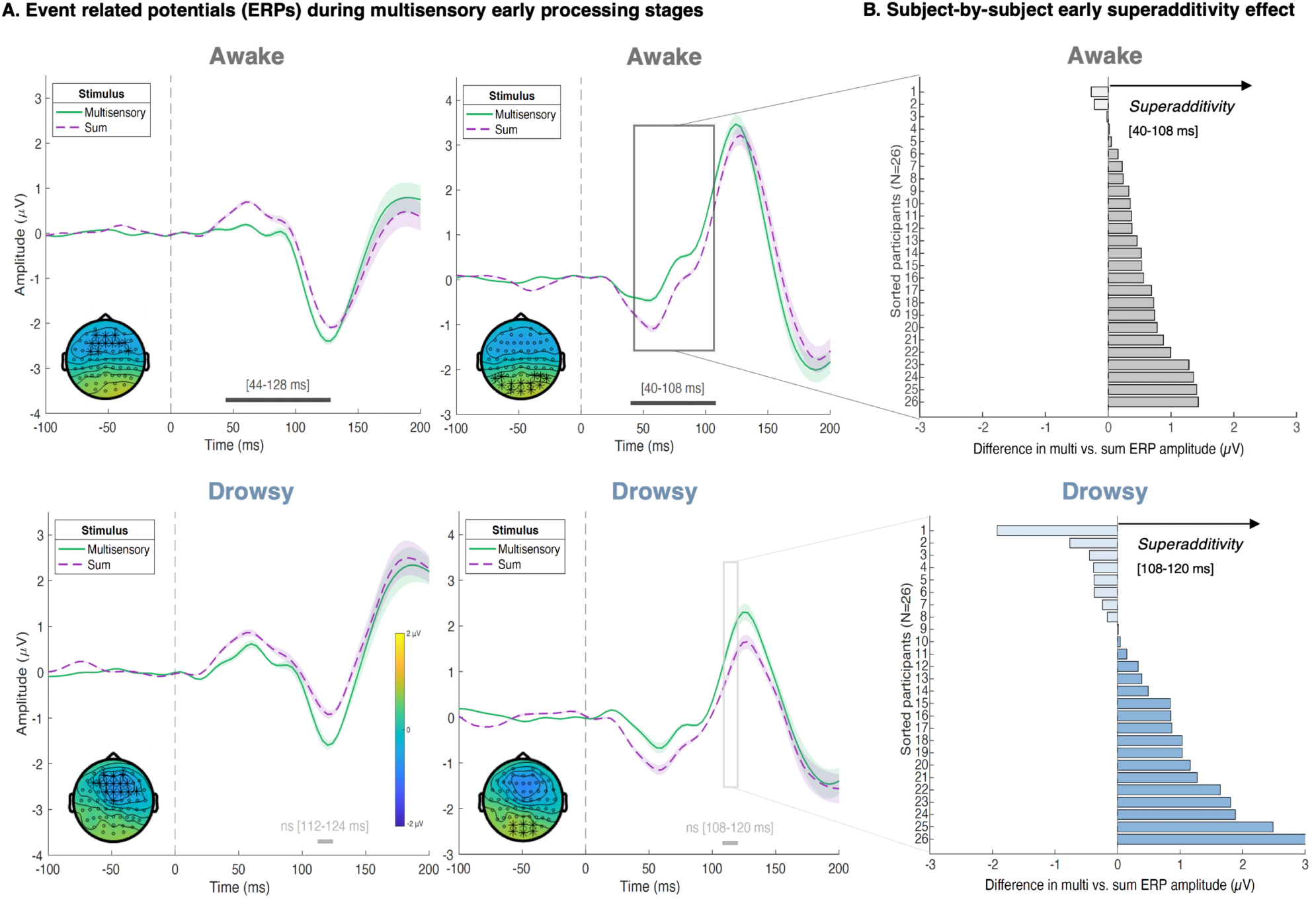
Early event-related potential (ERP) elicited by multisensory and sum of the unisensory conditions during wakefulness and drowsiness. The plots show the average ERP across all electrodes that were part of the observed cluster (indicated by black stars in the topoplots), where the shaded area corresponds to the standard error. The topoplots show the average difference between the ERPs evoked by the audiotactile and the sum of unisensory stimuli (auditory + tactile) for the time window indicated by the grey horizontal bars along the x-axis. A) During early sensory encoding (<150 ms post-stimulus), cluster-based permutation analyses revealed the classical multisensory superadditivity effect (higher ERP amplitude associated with responses to multisensory stimuli compared to the sum of unisensory responses) during wakefulness. This effect was not observed in drowsiness, where clusters (with comparisons indicated as non-significant [ns]) are shown for consistency and transparency, illustrating the pattern observed across participants. However, no significant between-state differences (multi-sum difference of differences) were observed before 150 ms, suggesting that although the effect appears state-dependent at encoding, statistical support for an early difference is lacking. B) Subject-by-subject mean voltage difference between multisensory - sum in wakefulness (grey bars) and drowsiness (blue bars). For the awake state, differences for each subject were calculated across the posterior electrodes included in the significant positive cluster shown in the panel immediately to the left, during the 40–108 ms time window. For the drowsy state, differences were calculated across the electrodes forming the detected (although not statistically significant) cluster displayed in the left panel, during the 108–120 ms time window. Difference in ERP amplitude (sum - multi) was computed after adding a constant positive offset to all values prior to averaging, ensuring polarity did not affect the estimate. Darker coloured lines highlight subjects showing effects in the direction of the superadditivity effect.

Late processing stages analyses, commonly related to decision making and motor response execution phase, showed an opposite pattern. We observed a reliable lower voltage amplitude in the multisensory condition compared to the sum of the two unisensory conditions in the same frontal-central and parietal-occipital electrodes from ∼200 ms onwards in the awake condition (frontal: 200-500 ms, *p* < 0.001, increased negativity for unisensory vs. multisensory; parietal: 192-500 ms, *p* < 0.001, increased positivity). This same pattern (i.e., significant differences in ERP amplitude between multisensory vs. sum of the unisensory) was also observed in the drowsy condition, although ∼100 ms delayed compared to wakefulness (frontal: 288-500 ms, *p* < 0.001; increased negativity for unisensory vs. multisensory y; parietal: 290-500 ms, *p* < 0.001; increased positivity), see Figure 4A. This pattern, referred to as subadditivity in the additive model framework, has been previously reported as a multisensory integration marker during late processing stages in similar audio–tactile detection tasks (Murray et al., 2005). A direct comparison between awake and drowsy conditions (multi–sum difference of differences) further revealed that this subadditivity effect was significantly reduced during drowsiness in both positive (160–388 ms) and negative (192–420 ms) electrode clusters (see Figure 4A-B; clusters of the awake vs. drowsy difference are represented in Supplemental Fig. S5). Notably, no differences in peak latency were observed between wakefulness and drowsiness, indicating that the cluster effects reflect a delay in the emergence of statistically reliable differences between conditions, rather than shifts in ERP peak latency (see Text S1).

**Figure 4.**
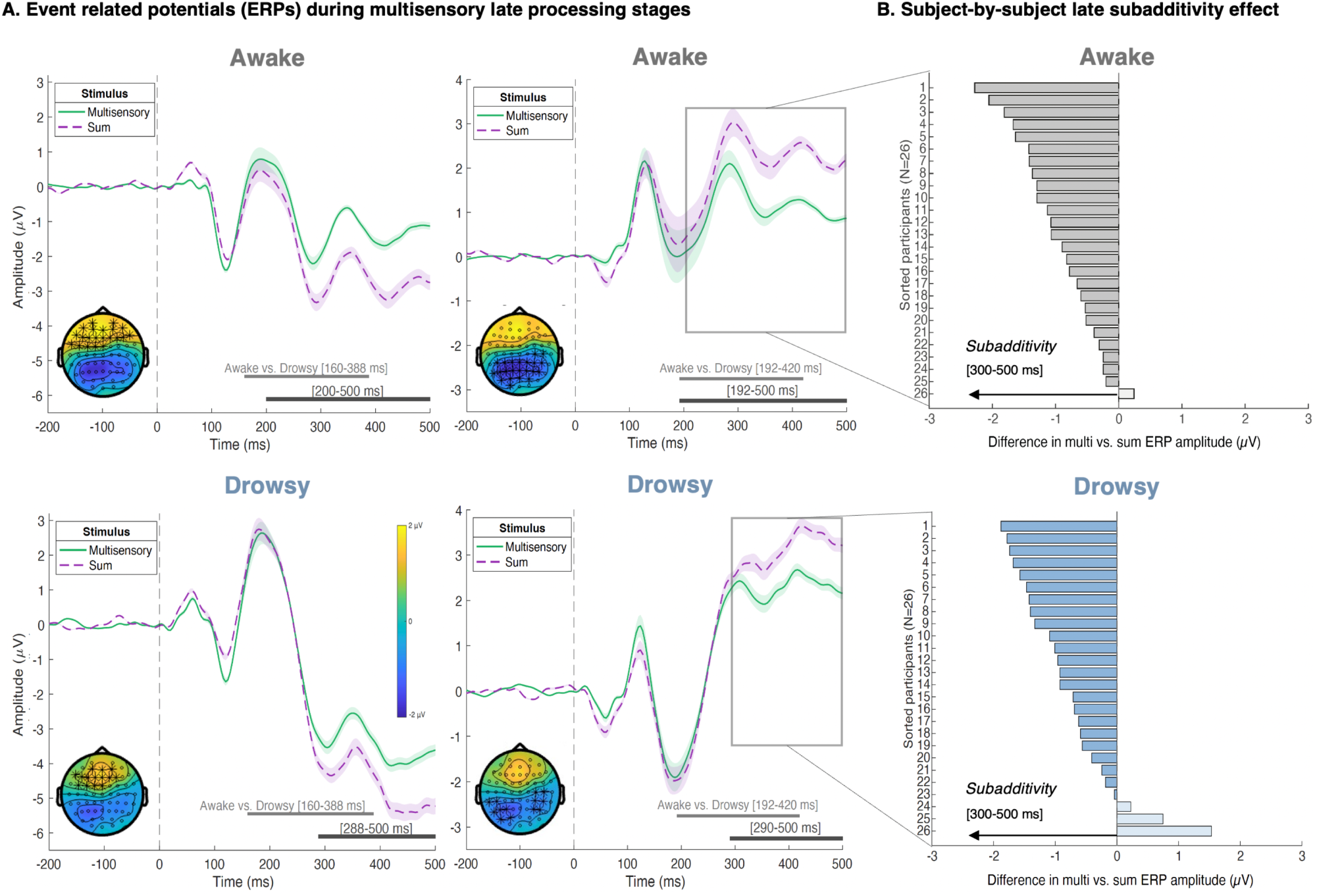
Late ERPs elicited by multisensory and sum of the unisensory conditions during wakefulness and drowsiness. A) During late decision-making and response execution processing stages (>200 ms post-stimulus), an opposite subadditivity pattern (reduced ERP amplitude in the multisensory condition compared to the summed unisensory responses) emerged in the same two electrodes’ areas (frontal and parietal) from ∼200 ms onward during wakefulness, and at ∼300 ms when drowsy. A direct comparison between awake and drowsy conditions (multi–sum difference of differences) further revealed that this subadditivity effect was significantly reduced during drowsiness, as depicted by the light grey horizontal line. B) Subject-by-subject mean voltage difference between multisensory - sum in wakefulness (grey bars) and drowsiness (blue bars) across the posterior electrodes involved in each corresponding detected positive cluster and during each cluster’s latency. Darker coloured lines highlight subjects showing effects in the direction of the subadditivity effect.

### Multisensory neural representations remain detectable but are less temporally stable during drowsiness

Our hypothesis-driven ERP analysis, targeting early markers of multisensory integration (the classical superadditivity effect), showed robust evidence of this effect during wakefulness, whereas it was not consistently observed under drowsiness. To examine whether a more spatially or temporally extended pattern of neural activity might support multisensory integration in the drowsy state, we conducted a multivariate decoding approach.

We trained linear discriminant analysis (LDA) classifiers to decode multisensory vs. the sum of unisensory responses, achieving above-chance decoding during both early and late processing stages in wakefulness and drowsiness (∼90-500 ms; see Figure 5A). Temporal generalization analyses revealed a broad generalization of neural representations associated with multisensory interactions from shortly after ∼100 ms onwards in both states. As expected, these representations seemed to be more stable and sustained during wakefulness compared to drowsiness (see Figure 5B).

**Figure 5.**
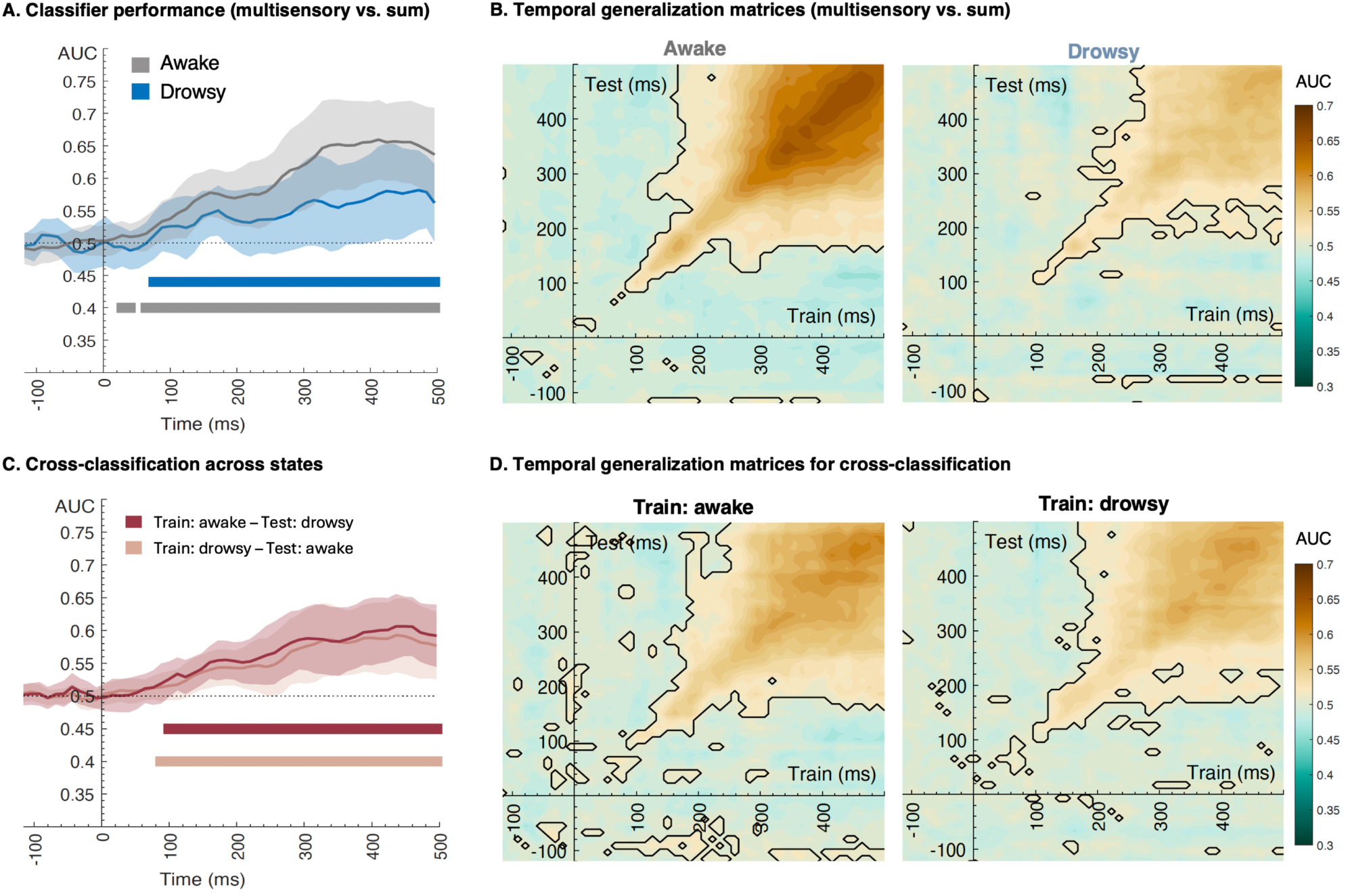
Multivariate classification of stimuli across alertness states. A) LDA classifier performance decoding type of stimulus (multisensory vs. sum of unisensory). Both awake (grey) and drowsy (blue) states showed above-chance decoding during early and late processing stages. Shaded areas represent standard error. Horizontal bars denote significant differences between stimuli conditions (cluster-corrected, *p* < 0.05). B) Temporal generalization matrices of decoding performance (multisensory vs. sum of unisensory responses) for wakefulness (left) and drowsiness (right). Both states showed significant decoding from ∼100 ms onwards, although decoding accuracy was higher and more sustained in wakefulness. Generalization across time indicates stable neural representations supporting multisensory interactions. Black contours denote clusters of significance (cluster-corrected, p < 0.05). C) Classifiers trained in wakefulness and tested in drowsiness (dark red) and vice versa (light red) showed above-chance decoding of multisensory versus unisensory-sum conditions within ∼100–500 ms, indicating similar patterns of neural activity across states. D) Temporal generalization matrices further confirmed stable cross-state representations, with classifiers trained at specific time points generalizing broadly across subsequent time windows.

### Shared multisensory representations generalize across alertness states early in processing

We further examined whether multisensory interactions are represented similarly across levels of alertness to determine whether the spatiotemporal patterns of brain activity that account for the differences between the multisensory condition and the summed condition are comparable during wakefulness and drowsiness. Cross-classification across states and stimulus conditions revealed similar neural patterns in wakefulness and drowsiness, supporting common neural representations of relevant multisensory information within ∼100–500 ms (see Figure 5C). Moreover, the temporal generalization analysis indicated that these cross-state representations were not only detectable but also temporally stable (see Figure 5D).

## DISCUSSION

Fluctuations in alertness levels occur dynamically and nonlinearly throughout the day, reflecting gradual shifts along the wake–sleep continuum. Accumulating evidence indicates that these fluctuations modulate the way we process and respond to unisensory events (for a review, see Lacaux et al., 2024). By contrast, its impact on multisensory integration, a fundamental process through which the brain combines signals across modalities to support adaptive behavior, remains poorly understood. Here, we revealed that the classic redundancy effect was largely preserved during drowsiness, albeit overall responses were slower. Analyses of full RT further showed that empirical multisensory benefits and violations of Miller’s bound (a hallmark of multisensory integration) were preserved during drowsiness, with no reliable differences across alertness levels. Moreover, the computational modelling framework suggests that these results cannot be explained by general response slowing, sequential dependencies or changes in unisensory response dynamics (Couth et al., 2018; Mahoney & Verghese, 2022). Together, behavioural and computational findings indicate that redundancy benefits persist during the transition into sleep, partially aligning with previous evidence. While prior work has suggested that multisensory signals may offset behavioural costs of reduced alertness (Artigas et al., 2025), our results show that these benefits do not fully compensate for the overall slowing observed during drowsiness.

Despite this behavioural preservation, a different picture emerged in the neural domain. During wakefulness, ERP analyses revealed two additive model-based signatures traditionally associated with multisensory integration: early superadditivity within 150 ms, and late subadditivity from ∼200 ms onwards (see Murray et al., 2005, for a similar result). By contrast, during drowsiness, early superadditivity was no longer detectable while late subadditivity was reduced in magnitude and delayed relative to wakefulness. Although this pattern aligns with previous evidence that canonical univariate signatures can diminish or even disappear during drowsiness despite preserved behaviour (e.g., Canales-Johnson et al., 2020), direct between-state comparisons did not yield reliable early differences. This may reflect substantial inter-individual variability, but it may also indicate that meta-stable state-related differences are subtle rather than qualitative. Consistent with this interpretation, ERPs spatiotemporal dynamics were comparable across states, and the heightened trial-to-trial variability characteristic of low alertness (Andrillon et al., 2021; Comsa et al., 2019; Strauss et al., 2015) may contribute to a reduced signal-to-noise ratio, potentially limiting the detectability of subtle univariate effects. Future work will be needed to directly characterise how transient fluctuations within drowsiness relate to neural stability.

One might speculate that the early superadditivity observed in wakefulness could reflect the principle of inverse effectiveness, which predicts stronger multisensory enhancement when unisensory inputs are weak (Meredith & Stein, 1983; Stein & Stanford, 2008). Yet, if this principle held in humans, greater enhancement should emerge under drowsiness, when perceptual thresholds rise (e.g., Goupil & Bekinschtein, 2012; Ogilvie & Wilkinson, 1984). Instead, we observed reduced early multisensory effects, suggesting that inverse effectiveness alone cannot account for our findings. This interpretation is consistent with evidence indicating that, in humans, such effects are highly context-dependent and may not generalise beyond tightly controlled or animal-based paradigms (Holmes, 2007; Stevenson et al., 2012). Regarding the later subadditive effects, caution is warranted in interpreting them, as they may reflect also shared, non-sensory activity (e.g., motor preparation) that is counted once in multisensory responses but summed across unisensory conditions (Besle et al., 2004, 2009). Thus, although our results align with prior reports of late subadditivity in audio-tactile paradigms, this component might be reflecting a mixture of multisensory interactions and overlapping non-sensory processes.

A clearer picture emerged when considering distributed neural patterns. Robust multisensory decoding was evident in both alertness regimes, emerging from ∼80 ms and extending into late processing periods. However, neural representations seemed more sustained and temporally coherent during wakefulness and decayed more rapidly during drowsiness, consistent with reduced temporal stability under low alertness (Canales-Johnson et al., 2020; Jagannathan et al., 2022; Noreika et al., 2020). Cross-states decoding however revealed shared spatiotemporal patterns from ∼100 ms onwards, suggesting that although the temporal stability of these representations can be modulated by alertness, their underlying structure remains largely preserved. Thus, drowsiness may weaken but not qualitatively alter multisensory integration. That is, even if a doctor’s reaction to the pager alert slows under drowsiness, the benefit of receiving simultaneous auditory and tactile cues is likely to persist, supported by neural mechanisms similar to those observed during wakefulness.

Despite the long-standing view that multisensory integration constitutes a robust property of the nervous system, even shown to persist at the single-cell level under anaesthesia (e.g., Alais et al., 2010; Deroy et al., 2014; Meredith & Stein, 1983), our results also highlight substantial variability across individuals. Not all participants exhibited redundancy gains or violated Miller’s bound, even in wakefulness (19 out of 26), and equivalent variability was observed in neural signatures such as superadditivity. While imperfect adherence to speeded-response instructions may explain part of this variance, the pattern suggests that multisensory integration is not a fixed function, but a conserved computation expressed with meaningful variability across individuals and conditions. Consistent with this view, converging evidence indicates that multisensory processing is dynamically shaped by ongoing brain states and task demands. Pre-stimulus power and phase of oscillatory dynamics in alpha and beta bands have been shown to modulate multisensory interactions and bias whether signals from different sensory modalities are perceived as integrated or segregated (e.g., Keil et al., 2012, 2014; Kaiser et al., 2019; Leonardelli et al., 2015). In parallel, cognitive factors such as attentional load have been shown to alter multisensory integration (Michail & Keil, 2018; Michail et al., 2021), further challenging the notion of an automatic process. More broadly, similar state-dependent mechanisms have been implicated across perceptual domains, where pre-stimulus oscillatory dynamics shape detection and perceptual outcome (e.g., Ai & Ro, 2014; Baumgarten et al., 2014; Strauss et al., 2022), and across more stable changes in brain function, including healthy ageing and neurological conditions (de Dieuleveult et al., 2017; Stevenson et al., 2014).

Within this broader framework, our findings extend previous work by showing that spontaneous fluctuations along the wake-sleep continuum, indexed here through the pre-stimulus theta/alpha ratio, modulate the neural expression of multisensory processing while leaving behavioural and computational indices largely preserved. This pattern may reflect, albeit speculatively, an intermediate point along a broader continuum, where multisensory computations are robustly expressed during wakefulness, remain functionally preserved but neurally altered during drowsiness, and may progressively degrade in deeper or unconscious sleep states. Consistent with this view, recent work on audiotactile peripersonal space processing has shown that a high-beta centro-parietal index of multisensory body-centred representation is present during wakefulness and dreaming, but absent during dreamless sleep (Serino et al., 2024). Although this marker differs from the detection-based indices used here, together these findings suggest that distinct forms of multisensory integration can exhibit graded sensitivity to changes in brain state and conscious level.

Notably, the percentile-based classification used here discretises what is inherently a continuous process. Accordingly, the “awake” and “drowsy” labels should be interpreted as reflecting metastable regimes (i.e., sets of microstates) positioned closer to the extremes of the alertness continuum, namely attentive wakefulness and sleep onset, respectively. Consistent with this, we observed differences in response speed within the “awake” category as a function of block context, despite comparable response rates, suggesting residual variability within these regimes. Future work will be needed to better characterise these within-state fluctuations and to explore alternative approaches for capturing alertness dynamics beyond percentile-based binning.

Beyond its theoretical significance, understanding how fluctuating alertness modulates multisensory integration has clear practical implications. Previous driving-simulator studies in fully alert participants have shown that audio–tactile warning signals can significantly speed up braking responses compared with unisensory cues (Ho et al., 2007), and that initial contact with rumble strips reduces physiological sleepiness as indexed by the Karolinska Drowsiness Scale (Watling et al., 2016). While these findings highlight the behavioural benefits of multisensory stimulation, they do not address how such benefits are mirrored in the brain as alertness declines. Our study extends previous works by providing temporally resolved neural evidence that the mechanisms supporting multisensory integration remain largely preserved during drowsiness, while contributing to the ongoing debate on where multisensory signals enter perceptual decision making, clarifying whether they arise during sensory encoding (e.g., Foxe & Schroeder, 2005) or emerge later during evidence accumulation and decision formation (e.g., Noppeney et al., 2010; Raposo et al., 2014).

In sum, our findings show that the core computations underlying multisensory integration remain largely preserved across the wake-sleep transition at the behavioural and computational level, despite alterations in their neural expression. This resilience likely reflects the behavioural importance of combining information across senses to guide action under fluctuating internal meta-stable states. Yet, the marked inter-individual variability observed here, together with the partial dissociation across behavioural, computational and neural indices, adds to growing evidence that multisensory integration is not uniformly expressed, but can vary as a function of ongoing brain states. As daily life regularly places humans in conditions of reduced alertness (from clinical night shifts to monotonous driving or everyday fatigue) understanding how the brain maintains multisensory processing during these transitions is essential.

## Data and code availability statement

The experimental protocol, hypotheses, and analysis plan were preregistered on OSF prior to data collection. All deviations from the preregistered protocol are transparently reported in the manuscript. Raw EEG data, organized according to the BIDS standard, are available on Zenodo (https://doi.org/10.5281/zenodo.17951706). Sorted data and the analysis-specific code used in this study are publicly available in the OSF repository (https://osf.io/7pt86/overview).

## Acknowledgements

This research was supported by a predoctoral fellowship from the Spanish Ministry of Science, Innovation and Universities to the first author (FPU21/00388). Luis F. Ciria was supported by project PID2022-142737NA-I00 funded by MCIN/AEI/10.13039/501100011033 and by FEDER, EU, and by Grant RYC2023-045268-I funded by MCIN/AEI/10.13039/501100011033 and by the European Union NextGenerationEU/PRTR. This research was also supported by Core Funds from the CCC-Lab. We thank Peter Macko (University of Granada) for his invaluable technical support in designing and implementing the experimental setup, including the custom hardware and software that enabled precise delivery of audio-tactile stimulation and accurate response recordings. We also thank Louis Roberts, Andres Canales-Johnson (University of Cambridge) and Marina Dauphin (Universidad de Granada) for their exploratory attempts to apply information-theoretic analyses to our data. Portions of the manuscript were edited with the assistance of an AI-based language model. The authors take full responsibility for the content.

## Author contributions

C.AL., D.S., T.A.B., and L.F.C. designed research; C.AL. and C.AV. performed research; C.AL. and C.AV. analyzed data; C.AL., D.S., and L.F.C. wrote the paper; All authors edited the manuscript and approved the final version; D.S., T.A.B., and L.F.C. supervised the project.

## REFERENCES

Ai, L., & Ro, T. (2014). The phase of prestimulus alpha oscillations affects tactile perception. Journal of Neurophysiology, 111(6), 1300–1307. 10.1152/jn.00125.2013

Alais, D., Newell, F., & Mamassian, P. (2010). Multisensory Processing in Review: From Physiology to Behaviour. Seeing and Perceiving, 23(1), 3–38. 10.1163/187847510X488603

Alameda, C., Avancini, C., Sanabria, D., Bekinschtein, T. A., Canales-Johnson, A., & Ciria, L. F. (2024). Staying in control: Characterizing the mechanisms underlying cognitive control in high and low arousal states. British Journal of Psychology (London, England: 1953). 10.1111/bjop.12715

Andrillon, T., Burns, A., Mackay, T., Windt, J., & Tsuchiya, N. (2021). Predicting lapses of attention with sleep-like slow waves. Nature Communications, 12(1), 3657. 10.1038/s41467-021-23890-7

Artigas, C., Morales-Torres, R., Rojas-Thomas, F., Villena-González, M., Rubio, I., Ramírez-Benavides, D., Bekinschtein, T., Campos-Arteaga, G., & Rodríguez, E. (2025). When alertness fades: Drowsiness-induced visual dominance and oscillatory recalibration in audiovisual integration. International Journal of Psychophysiology, 212, 112562. 10.1016/j.ijpsycho.2025.112562

Bates, D., Mächler, M., Bolker, B., & Walker, S. (2014). Fitting Linear Mixed-Effects Models using lme4 (arXiv:1406.5823). 10.48550/arXiv.1406.5823

Baumgarten, T., Schnitzler, A., & Lange, J. (2014). Prestimulus Alpha Power Influences Tactile Temporal Perceptual Discrimination and Confidence in Decisions. Cerebral cortex, 26. 10.1093/cercor/bhu247

Berman, A. L. (1961). Interaction of cortical responses to somatic and auditory stimuli in anterior ectosylvian gyrus of cat. Journal of Neurophysiology, 24(6), 608–620. 10.1152/jn.1961.24.6.608

Bertoni, T., Ricci, G., Jöhr, J., Donno, B., Fellrath, J., Stephan, A., Foglia, C., Lecci, S., Da Silva, M. L., Galigani, M., Pozeg, P., Dunet, V., Noël, J.-P., Magosso, E., Diserens, K., Siclari, F., & Serino, A. (2026). Multisensory integration in peripersonal space indexes consciousness states in sleep and disorders of consciousness. Cell Reports Medicine, 7(4), 102705. 10.1016/j.xcrm.2026.102705

Besle, J., Bertrand, O., & Giard, M.-H. (2009). Electrophysiological (EEG, sEEG, MEG) evidence for multiple audiovisual interactions in the human auditory cortex. Hearing Research, 258(1), 143–151. 10.1016/j.heares.2009.06.016

Besle, J., Fort, A., & Giard, M.-H. (2004). Interest and validity of the additive model in electrophysiological studies of multisensory interactions. Cognitive Processing, 5(3). 10.1007/s10339-004-0026-y

Canales-Johnson, A., Beerendonk, L., Blain, S., Kitaoka, S., Ezquerro-Nassar, A., Nuiten, S., Fahrenfort, J., van Gaal, S., & Bekinschtein, T. A. (2020). Decreased Alertness Reconfigures Cognitive Control Networks. The Journal of Neuroscience, 40(37), 7142–7154. 10.1523/JNEUROSCI.0343-20.2020

Ciria, L. F., Suárez-Pinilla, M., Williams, A. G., Jagannathan, S. R., Sanabria, D., & Bekinschtein, T. A. (2021). Different underlying mechanisms for high and low arousal in probabilistic learning in humans. Cortex, 143, 180–194. 10.1016/j.cortex.2021.07.002

Comsa, I. M., Bekinschtein, T. A., & Chennu, S. (2019). Transient Topographical Dynamics of the Electroencephalogram Predict Brain Connectivity and Behavioural Responsiveness During Drowsiness. Brain Topography, 32(2), 315–331. 10.1007/s10548-018-0689-9

Couth, S., Gowen, E., & Poliakoff, E. (2018). Using Race Model Violation to Explore Multisensory Responses in Older Adults: Enhanced Multisensory Integration or Slower Unisensory Processing? Multisensory Research, 31(3-4), 151–174. 10.1163/22134808-00002588

de Dieuleveult, A. L., Siemonsma, P. C., van Erp, J. B. F., & Brouwer, A.-M. (2017). Effects of Aging in Multisensory Integration: A Systematic Review. Frontiers in Aging Neuroscience, 9, 80. 10.3389/fnagi.2017.00080

De Gennaro, L., Ferrara, M., & Bertini, M. (2001). The boundary between wakefulness and sleep: Quantitative electroencephalographic changes during the sleep onset period. Neuroscience, 107(1), 1–11. 10.1016/S0306-4522(01)00309-8

Delorme, A., & Makeig, S. (2004). EEGLAB: An open source toolbox for analysis of single-trial EEG dynamics including independent component analysis. Journal of neuroscience methods, 134(1), 9–21.

Deroy, O., Chen, Y.-C., & Spence, C. (2014). Multisensory constraints on awareness. Philosophical Transactions of the Royal Society B: Biological Sciences, 369(1641), 20130207. 10.1098/rstb.2013.0207

Foxe, J. J., & Schroeder, C. E. (2005). The case for feedforward multisensory convergence during early cortical processing. Neuroreport, 16(5), 419–423. 10.1097/00001756-200504040-00001

Goupil, L., & Bekinschtein, T. (2012). Cognitive processing during the transition to sleep. Archives Italiennes de Biologie, 150(2/3), Article 2/3. 10.4449/aib.v150i2.1247

Grootswagers, T., Wardle, S. G., & Carlson, T. A. (2017). Decoding Dynamic Brain Patterns from Evoked Responses: A Tutorial on Multivariate Pattern Analysis Applied to Time Series Neuroimaging Data. Journal of Cognitive Neuroscience, 29(4), 677–697. 10.1162/jocn_a_01068

Helton, W. S., & Russell, P. N. (2015). Rest is best: The role of rest and task interruptions on vigilance. Cognition, 134, 165–173. 10.1016/j.cognition.2014.10.001

Ho, C., Reed, N., & Spence, C. (2007). Multisensory in-car warning signals for collision avoidance. Human Factors, 49(6), 1107–1114. 10.1518/001872007X249965

Holmes, N. P. (2007). The law of inverse effectiveness in neurons and behaviour: Multisensory integration versus normal variability. Neuropsychologia, 45(14), 3340–3345. 10.1016/j.neuropsychologia.2007.05.025

Innes, B. R., & Otto, T. U. (2019). A comparative analysis of response times shows that multisensory benefits and interactions are not equivalent. Scientific Reports, 9(1), 1–10.

Jagannathan, S. R., Bareham, C. A., & Bekinschtein, T. A. (2022). Decreasing Alertness Modulates Perceptual Decision-Making. Journal of Neuroscience, 42(3), 454–473. 10.1523/JNEUROSCI.0182-21.2021

Jagannathan, S. R., Ezquerro-Nassar, A., Jachs, B., Pustovaya, O. V., Bareham, C. A., & Bekinschtein, T. A. (2018). Tracking wakefulness as it fades: Micro-measures of alertness. NeuroImage, 176, 138–151. 10.1016/j.neuroimage.2018.04.046

Johns, M. W. (1991). A New Method for Measuring Daytime Sleepiness: The Epworth Sleepiness Scale. Sleep, 14(6), 540–545. 10.1093/sleep/14.6.540

Johns, M. W. (1992). Reliability and factor analysis of the Epworth Sleepiness Scale. Sleep: Journal of Sleep Research & Sleep Medicine, 15(4), 376–381. 10.1093/sleep/15.4.376

Kaiser, M., Senkowski, D., Busch, N. A., Balz, J., & Keil, J. (2019). Single trial prestimulus oscillations predict perception of the sound-induced flash illusion. Scientific Reports, 9(1), 5983. 10.1038/s41598-019-42380-x

Keil, J., Müller, N., Hartmann, T., & Weisz, N. (2014). Prestimulus beta power and phase synchrony influence the sound-induced flash illusion. Cerebral Cortex, 24(5), 1278–1288. 10.1093/cercor/bhs409

Keil, J., Müller, N., Ihssen, N., & Weisz, N. (2012). On the Variability of the McGurk Effect: Audiovisual Integration Depends on Prestimulus Brain States. Cerebral Cortex, 22(1), 221–231. 10.1093/cercor/bhr125

Kleiner, M., Brainard, D., & Pelli, D. (2007). What’s new in Psychtoolbox*-*3*?* 14. 10.1177/03010066070360S101

Kuznetsova, A., Brockhoff, P. B., & Christensen, R. H. B. (2017). lmerTest Package: Tests in Linear Mixed Effects Models. Journal of Statistical Software, 82, 1–26. 10.18637/jss.v082.i13

Lacaux, C., Strauss, M., Bekinschtein, T. A., & Oudiette, D. (2024). Embracing sleep-onset complexity. Trends in Neurosciences, 47(4), 273–288. 10.1016/j.tins.2024.02.002

Leonardelli, E., Braun, C., Weisz, N., Lithari, C., Occelli, V., & Zampini, M. (2015). Prestimulus oscillatory alpha power and connectivity patterns predispose perceptual integration of an audio and a tactile stimulus. Human Brain Mapping, 36(9), 3486–3498. 10.1002/hbm.22857

López-García, D., Peñalver, J. M., Górriz, J. M., & Ruz, M. (2022). MVPAlab: A machine learning decoding toolbox for multidimensional electroencephalography data. Computer Methods and Programs in Biomedicine, 214, 106549.

Maris, E., & Oostenveld, R. (2007). Nonparametric statistical testing of EEG-and MEG-data. Journal of neuroscience methods, 164(1), 177–190.

Mercier, M. R., & Cappe, C. (2020). The interplay between multisensory integration and perceptual decision making. NeuroImage, 222, 116970.

Meredith, M. A., & Stein, B. E. (1983). Interactions Among Converging Sensory Inputs in the Superior Colliculus. Science, 221(4608), 389–391. 10.1126/science.6867718

Miller, J. (1982). Divided attention: Evidence for coactivation with redundant signals. Cognitive Psychology, 14(2), 247–279. 10.1016/0010-0285(82)90010-X

Michail, G., & Keil, J. (2018). High cognitive load enhances the susceptibility to non-speech audiovisual illusions. Scientific Reports, 8(1), 11530. 10.1038/s41598-018-30007-6

Michail, G., Senkowski, D., Niedeggen, M., & Keil, J. (2021). Memory Load Alters Perception-Related Neural Oscillations during Multisensory Integration. The Journal of Neuroscience, 41(7), 1505–1515. 10.1523/JNEUROSCI.1397-20.2020

Murray, M. M., Molholm, S., Michel, C. M., Heslenfeld, D. J., Ritter, W., Javitt, D. C., Schroeder, C. E., & Foxe, J. J. (2005). Grabbing Your Ear: Rapid Auditory–Somatosensory Multisensory Interactions in Low-level Sensory Cortices Are Not Constrained by Stimulus Alignment. Cerebral Cortex, 15(7), 963–974. 10.1093/cercor/bhh197

Noppeney, U., Ostwald, D., & Werner, S. (2010). Perceptual Decisions Formed by Accumulation of Audiovisual Evidence in Prefrontal Cortex. The Journal of Neuroscience, 30(21), 7434–7446. 10.1523/JNEUROSCI.0455-10.2010

Noreika, V., Kamke, M. R., Canales-Johnson, A., Chennu, S., Bekinschtein, T. A., & Mattingley, J. B. (2020). Alertness fluctuations when performing a task modulate cortical evoked responses to transcranial magnetic stimulation. NeuroImage, 223, 117305. 10.1016/j.neuroimage.2020.117305

Ogilvie, R. D., & Wilkinson, R. T. (1984). The Detection of Sleep Onset: Behavioral and Physiological Convergence. Psychophysiology, 21(5), 510–520. 10.1111/j.1469-8986.1984.tb00234.x

Oostenveld, R., Fries, P., Maris, E., & Schoffelen, J.-M. (2011). FieldTrip: Open Source Software for Advanced Analysis of MEG, EEG, and Invasive Electrophysiological Data. Computational Intelligence and Neuroscience, 2011, 1–9. 10.1155/2011/156869

Quinn, B. T., Carlson, C., Doyle, W., Cash, S. S., Devinsky, O., Spence, C., Halgren, E., & Thesen, T. (2014). Intracranial Cortical Responses during Visual–Tactile Integration in Humans. Journal of Neuroscience, 34(1), 171–181. 10.1523/JNEUROSCI.0532-13.2014

Raposo, D., Kaufman, M. T., & Churchland, A. K. (2014). A category-free neural population supports evolving demands during decision-making. Nature Neuroscience, 17(12), 1784–1792. 10.1038/nn.3865

Ross, H. A., Russell, P. N., & Helton, W. S. (2014). Effects of breaks and goal switches on the vigilance decrement. Experimental Brain Research, 232(6), 1729–1737. 10.1007/s00221-014-3865-5

Saper, C. B., Fuller, P. M., Pedersen, N. P., Lu, J., & Scammell, T. E. (2010). Sleep State Switching. Neuron, 68(6), 1023–1042. 10.1016/j.neuron.2010.11.032

Schumann, F., Steinborn, M. B., Kürten, J., Cao, L., Händel, B. F., & Huestegge, L. (2022). Restoration of Attention by Rest in a Multitasking World: Theory, Methodology, and Empirical Evidence. Frontiers in Psychology, 13. 10.3389/fpsyg.2022.867978

Shaw, L. H., Freedman, E. G., Crosse, M. J., Nicholas, E., Chen, A. M., Braiman, M. S., Molholm, S., & Foxe, J. J. (2020). Operating in a Multisensory Context: Assessing the Interplay Between Multisensory Reaction Time Facilitation and Inter-sensory Task-switching Effects. Neuroscience, 436, 122–135. 10.1016/j.neuroscience.2020.04.013

Stein, B. E., & Stanford, T. R. (2008). Multisensory integration: Current issues from the perspective of the single neuron. Nature reviews neuroscience, 9(4), 255–266.

Stevenson, R. A., Bushmakin, M., Kim, S., Wallace, M. T., Puce, A., & James, T. W. (2012). Inverse Effectiveness and Multisensory Interactions in Visual Event-Related Potentials with Audiovisual Speech. Brain topography, 25(3), 308–326. 10.1007/s10548-012-0220-7

Stevenson, R. A., Ghose, D., Fister, J. K., Sarko, D. K., Altieri, N. A., Nidiffer, A. R., Kurela, L. R., Siemann, J. K., James, T. W., & Wallace, M. T. (2014). Identifying and Quantifying Multisensory Integration: A Tutorial Review. Brain Topography, 27(6), 707–730. 10.1007/s10548-014-0365-7

Strauss, M., Sitt, J. D., King, J.-R., Elbaz, M., Azizi, L., Buiatti, M., Naccache, L., Van Wassenhove, V., & Dehaene, S. (2015). Disruption of hierarchical predictive coding during sleep. Proceedings of the National Academy of Sciences of the United States of America, 112(11), E1353–E1362. Scopus. 10.1073/pnas.1501026112

Strauss, M., Sitt, J. D., Naccache, L., & Raimondo, F. (2022). Predicting the loss of responsiveness when falling asleep in humans. NeuroImage, 251, 119003. 10.1016/j.neuroimage.2022.119003

Watling, C. N., Åkerstedt, T., Kecklund, G., & Anund, A. (2016). Do repeated rumble strip hits improve driver alertness? Journal of Sleep Research, 25(2), 241–247. 10.1111/jsr.12359

Xu, Y., Wokke, M., Noreika, V., Bareham, C., Jagannathan, S., Georgieva, S., Trentin, C., & Bekinschtein, T. (2025). Effects of alertness on perceptual detection and discrimination. Cortex, 190, 262–285. 10.1016/j.cortex.2025.06.018

