## Supplemental Text S1, Figure S1, S2, S3, S4, S5 for "Drowsiness alters the neural dynamics but not the core computations of multisensory integration"

### <u>Supplemental Material</u>

#### (Text S1)

***Control analyses for alertness classification based on the theta/alpha ratio***

For each participant, we present representative examples of the temporal evolution of the prestimulus theta/alpha ratio across the experimental session (Supplemental Figure S1), following the same format as Fig. 1C in the main manuscript (see Methods section "Alertness classification"). Complete participant-level plots for all participants are available in the study OSF repository (Theta-alpha_ratio_plots_all_subjects.pdf; https://osf.io/7pt86/files/osfstorage). Each panel displays single-trial data across the session (x-axis: pre-trial periods). The black trace represents reaction times (left y-axis), while blue markers indicate the theta/alpha ratio computed for each trial (right y-axis). Trials without responses (omissions) are shown as red markers. Vertical dashed lines denote transitions between experimental blocks, with awake and drowsy blocks labelled along the x-axis.

Across participants, these plots reveal gradual and within-session fluctuations in alertness. Periods characterised by lower theta/alpha values tend to coincide with faster and more stable reaction times and a low incidence of omissions, consistent with relatively higher levels of alertness. Conversely, increases in the theta/alpha ratio are typically accompanied by slower and more variable responses, as well as a higher frequency of omissions, reflecting reduced alertness. Overall, these patterns illustrate the continuous and dynamic nature of alertness fluctuations along the wake–sleep continuum.

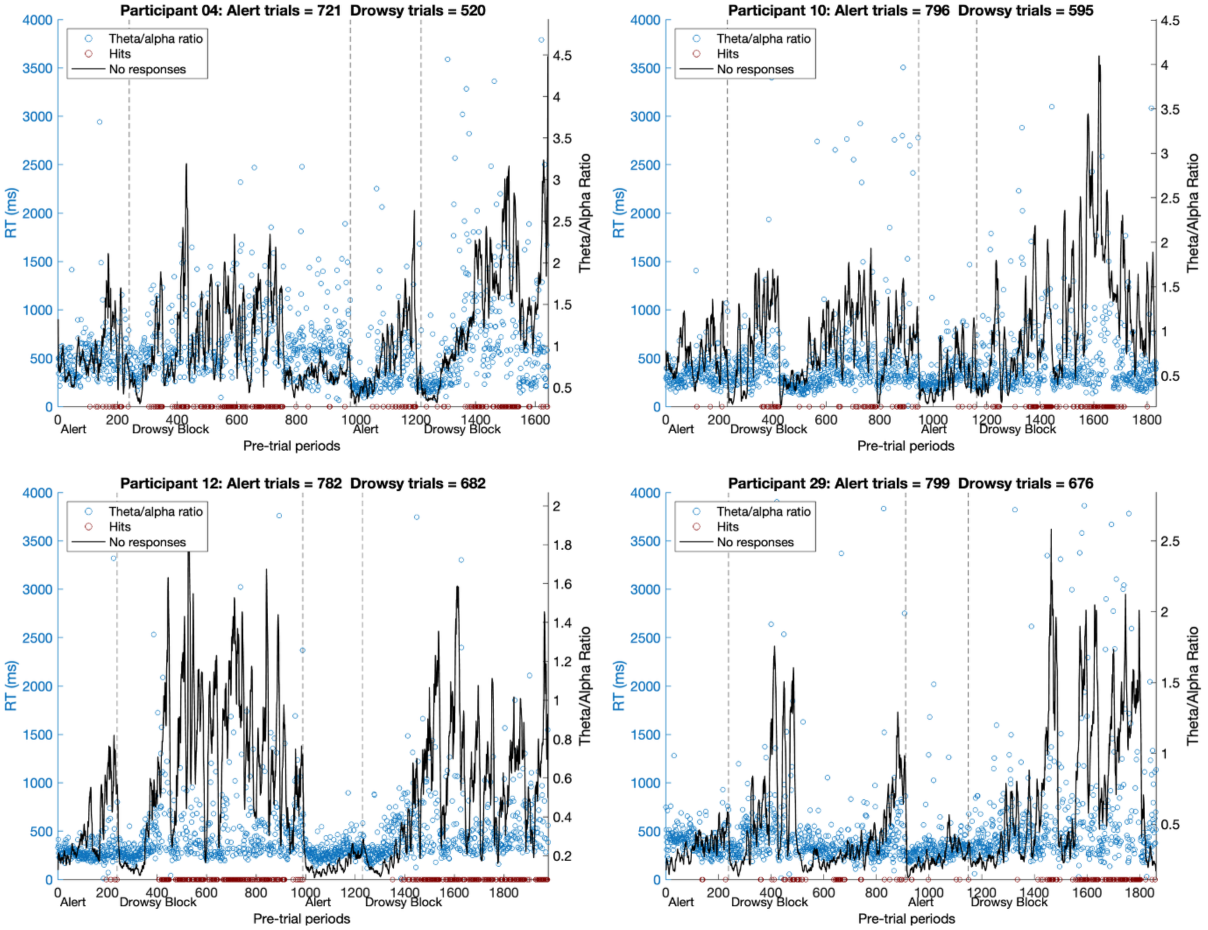


**Supplemental Figure S1.** Subject-level dynamics of alertness across the experimental session. For each participant, we display the full temporal evolution of the prestimulus theta/alpha ratio and behavioural markers (see description above). Panel titles indicate the participant identifier and the number of trials classified as awake and drowsy following the alertness classification procedure described in the Methods section.

During peer review, concerns were raised regarding the potential impact of individual differences in spectral properties (specifically, variability in alpha peak frequency and the contribution of the aperiodic component) on the classification of alertness states. To address this, we conducted a series of control analyses to assess the robustness of our classification procedure.

First, we recomputed the alertness index using individualized spectral parameters. For each participant, the aperiodic component of the EEG power spectrum was separated, and the individual alpha peak frequency was identified within the 8–13 Hz range. Based on this estimate, a personalized alpha band was defined as ±1 Hz around the peak, and the theta/alpha ratio was recomputed on a per-trial basis using these individualized parameters. We then compared the resulting trial classifications with those obtained using the original canonical bands (theta: 4–6 Hz; alpha: 10–12 Hz). As shown in Supplemental Fig. S2 (above), the overlap between the two classification approaches was high, ranging from approximately 70% to 90% across participants and alertness states. This indicates substantial consistency between methods despite differences in spectral parametrization.

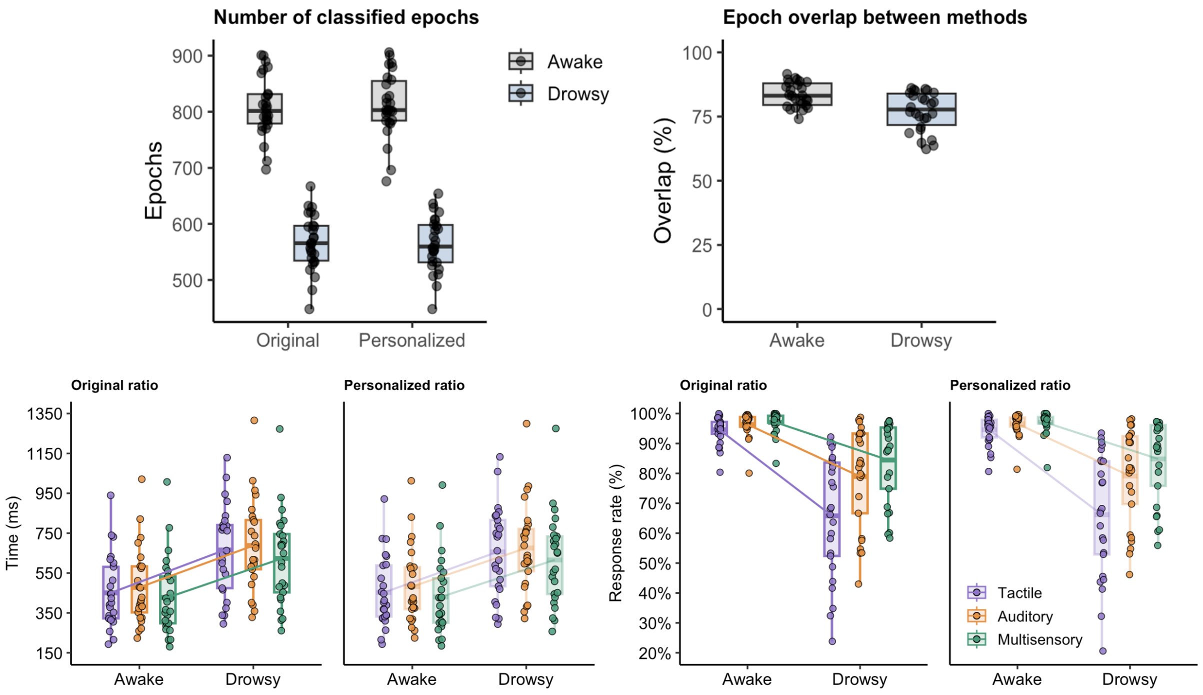


**Supplemental Figure S2.** Above, consistency of alertness classification across methods. Left panel shows the number of trials classified as awake and drowsy using the original and personalized theta/alpha ratio. Right panel shows the percentage of overlap between both classification methods for awake and drowsy states. In all panels, each dot represents one participant. Boxplots display the distribution across participants, with the central line indicating the median, the box representing the interquartile range (IQR), and whiskers extending to 1.5× IQR. Below, behavioural results using original and personalized alertness classification. Reaction times (left panels) and response rates (right panels) are shown for each stimulus condition (tactile, auditory, multisensory) and alertness state (awake, drowsy), computed using the original (left) and personalized (right) theta/alpha ratio. Each dot represents the mean value for one participant in each condition.

To further assess the impact of this alternative classification on the main findings, we repeated the behavioural analyses using the personalized alertness labels. Reaction times (RTs) were analysed using a linear mixed-effects model including Method (original vs. personalized), State (awake vs. drowsy), and Stimulus type (tactile, auditory, multisensory) as fixed effects, with participant as a random factor. This analysis revealed no main effect of Method, $F(1,264) = 0.13$, $p = .72$, $\eta^2_p < 0.001$, and no interactions involving Method (Method × State: $F(1,264) = 0.42$, $p = 0.52$; Method × Stimulus: $F(2,264) < 0.01$, $p = 0.99$; Method × State × Stimulus: $F(2,264) = 0.01$, $p = 0.99$). In contrast, the expected main effects of State, $F(1,264) = 420.35$, $p < 0.001$, $\eta^2_p = 0.61$, and Stimulus, $F(2,264) = 11.27$, $p < 0.001$, $\eta^2_p = 0.08$, were preserved (see Supplemental Fig. S2, below left).

Equivalent analyses on response rate yielded the same pattern. There was no main effect of Method, $F(1,264) = 0.02$, $p = 0.88$, $\eta^2_p < 0.001$, nor any interactions involving Method (all *p*s > 0.90), while the main effects of State, $F(1,264) = 304.69$, $p < 0.001$, $\eta^2_p = 0.54$, and Stimulus, $F(2,264) = 31.49$, $p < 0.001$, $\eta^2_p = 0.19$, as well as their interaction, $F(2,264) = 18.42$, $p < 0.001$, $\eta^2_p = 0.12$, remained unchanged (see Supplemental Fig. S2, below right). Post hoc comparisons confirmed that there were no significant differences between the original and personalized classification methods in any condition for either RT or response rate (all Bonferroni-corrected *p*s > 0.60). Together, these results show that the classification of alertness is robust to individual differences in spectral features and are not driven by the specific choice of canonical frequency bands.

Following another suggestion raised during peer review, we conducted an additional behavioural analysis to examine whether awake trials differed depending on the block context in which they occurred. Specifically, we compared three trial types: (i) awake trials occurring within awake blocks, (ii) awake trials occurring within drowsy blocks, and (iii) drowsy trials.

RT analyses revealed a significant main effect of Block Context, $F(2,192) = 147.72$, $p < 0.001$, $\eta^2p = 0.61$, as well as a main effect of Stimulus, $F(2,192) = 5.53$, $p = 0.0046$, $\eta^2p = 0.05$, with no Block Context × Stimulus interaction ($F(4,192) = 0.09$, $p = 0.985$). Bonferroni-corrected post hoc comparisons showed that awake trials occurring within awake blocks were significantly faster than awake trials within drowsy blocks across all stimulus conditions (tactile: $p = 0.0002$; auditory: $p = 0.0001$; multisensory: $p = 0.0003$). Both categories of awake trials were, in turn, significantly faster than drowsy trials (all *p*s < 0.001; see Supplemental Figure S4, left). For response rate, the analysis revealed a main effect of Block Context, $F(2,192) = 119.51$, $p < 0.001$, $\eta^2p = 0.55$, a main effect of Stimulus, $F(2,192) = 13.72$, $p < 0.001$, $\eta^2p = 0.13$, and a significant Block Context × Stimulus interaction, $F(4,192) = 7.09$, $p < 0.001$, $\eta^2p = 0.13$. Importantly, Bonferroni-corrected post hoc comparisons showed no significant differences in response rate between awake trials from awake blocks and awake trials from drowsy blocks across all stimulus conditions (all *p*s ≥ 0.35). In contrast, both types of awake trials exhibited significantly higher response rates than drowsy trials (all *p*s < 0.001; see Supplemental Figure S4, right).

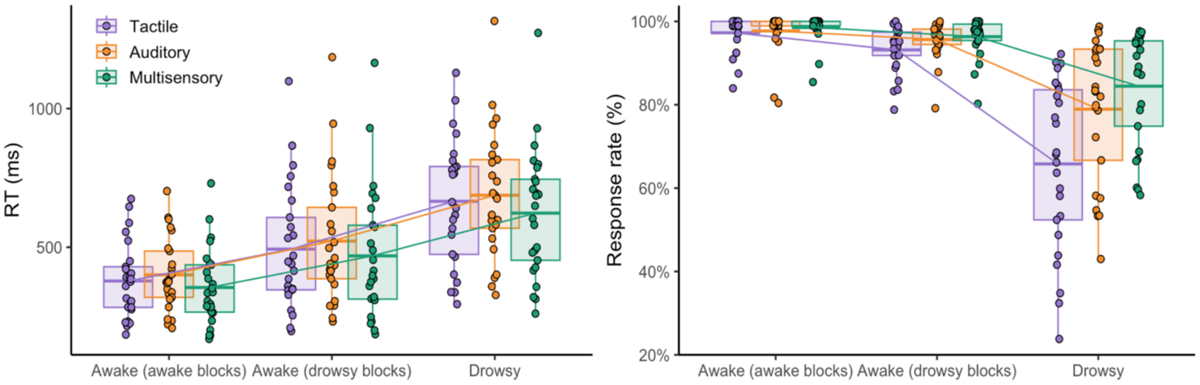


**Supplemental Figure S3.** Block context within the awake condition. Reaction times (left panel) and response rates (right panel) as a function of Block Context and Stimulus type. Three trial categories are shown: awake trials occurring in awake blocks, awake trials occurring in drowsy blocks, and drowsy trials. Boxplots represent the distribution of subject-level means for each condition. The central line indicates the median, the box spans the interquartile range (25th–75th percentiles), and whiskers extend to the most extreme values within 1.5 times the interquartile range. Each dot corresponds to the mean value for an individual participant in a given condition.

These results show that while RTs show a graded slowing for awake trials embedded within drowsy blocks, both categories of awake trials remain comparable in terms of behavioural responsiveness and clearly distinct from drowsy trials. This pattern is consistent with the view that alertness fluctuates continuously, in a dynamic and non-linear manner, and state categories here presented can be better understood as a regime of intermediate microstates (i.e., a metastable state) rather than discrete and fixed brain states, as previously reported in studies of cognitive dynamics during drowsiness (Canales-Johnson et al., 2020; Jagannathan et al., 2018; Lacaux et al., 2024).

#### *AIC-based selection of linear mixed-effects models*

To investigate whether the magnitude of multisensory temporal benefits (i.e., RSE) is modulated during the wake–sleep transition, we fitted RTs and response rate data using linear mixed-effects modelling, as implemented in the *lme4* R package (Bates et al., 2014). Akaike's Information Criterion (AIC) was used to compare competing model structures for each dependent variable. For RT, the model including the Alertness × Stimuli interaction provided the best fit (AIC = 3767.12), outperforming both the full model (Alertness × Stimuli × Side; AIC = 3823.04) and simpler alternatives, including models with Alertness only (AIC = 3809.76), Stimuli only (AIC = 4042.13), or a null model (AIC = 4061.82). In contrast, for response rate (ACC), the full model including Alertness × Stimuli × Side yielded the best fit (AIC = −470.22), outperforming reduced models including Alertness × Stimuli (AIC = −443.34), Alertness only (AIC = −385.02), Stimuli only (AIC = −233.96), and the null model (AIC = −221.83). Therefore, we selected the Alertness × Stimuli model for RT and the full Alertness × Stimuli × Side model for ACC for subsequent inference.

#### ***Linear mixed-effects analyses on log-transformed reaction times***

During the peer-review process, concerns were raised regarding the positive skew typically observed in RT distributions and its potential impact on linear mixed-effects modelling. To address this, we repeated the main RT analyses using log-transformed RTs. Specifically, RTs were log-transformed at the trial level and fitted using the same linear mixed-effects model structure as in the main analysis (Alertness × Stimuli, with random intercepts for participants). Model estimation and inference were conducted using the same procedures as for the raw RT data. In addition, post hoc pairwise comparisons were computed using estimated marginal means (*emmeans*) with Bonferroni correction, and model assumptions were evaluated through inspection of residuals and distributional properties.

Analyses on log-transformed RTs yielded the same pattern of results as those obtained with raw RTs. The linear mixed-effects model revealed significant main effects of Alertness, $F(1, 281) = 522.47$, $p < 0.001$, and Stimuli, $F(2, 281) = 16.14$, $p < 0.001$, with no significant interaction between factors, $F(2, 281) = 0.22$, $p = 0.80$. Bonferroni-corrected post hoc contrasts showed significantly faster responses for multisensory compared to auditory stimuli, $t(281) = 5.66$, $p < 0.001$, and compared to tactile stimuli, $t(281) = 3.29$, $p = 0.004$, whereas the difference between the two unisensory conditions did not reach significance, $t(281) = -2.37$, $p = 0.055$. Overall, these results closely replicate the pattern observed with raw RTs, confirming that the reported effects are robust to deviations from normality.

#### ***Robustness of race model estimates***

Previous work has suggested that race model violations can be influenced by trial-history or task-switching effects, particularly when successive trials differ in modality or response requirements (Innes & Otto, 2019; Shaw et al., 2020). To assess whether our estimates of Miller's bound violation, and especially the comparison between wakefulness and drowsiness, could be explained by this type of sequential dependency rather than by multisensory processing per se, we conducted an additional control analysis focused on trial history. For each trial, we classified the relation to the immediately preceding trial into four categories: no-switch (same modality and same side), modality-only switch (different modality, same side), side-only switch (same modality, different side), and double switch (different modality and different side). Trials were classified separately within each alertness state, and mean RTs were computed for each switch category. For each participant and state, switch costs were then estimated as the difference between the mean RT of each switch category and the corresponding no-switch baseline.

We first examined whether switch costs differed as a function of alertness state and switch type using a linear mixed-effects model with State, SwitchType, and their interaction as fixed effects, and participant as a random intercept. This analysis revealed no main effect of State, $F(1,125) = 0.19$, $p = 0.667$, no main effect of SwitchType, $F(2,125) = 0.24$, $p = 0.790$, and no State × SwitchType interaction,

$F(2,125) = 0.76$, $p = 0.471$. Distributions of switch costs for each condition are shown in Supplemental Figure Se.

We next asked whether individual differences in the change in switch costs from wakefulness to drowsiness predicted the corresponding change in Miller's bound violation. To test this, we correlated the awake-to-drowsy change in modality-only, side-only, and double-switch costs with the awake-to-drowsy change in violation values. None of these correlations were significant (Spearman correlations, modality-only: $\rho = 0.216$, $p = 0.290$; side-only: $\rho = -0.019$, $p = 0.926$; double-switch: $\rho = -0.187$, $p = 0.361$). A multiple regression including all three switching measures likewise did not explain variance in the change in Miller's bound violation, $F(3, 22) = 0.96$, $p = 0.430$, $R^2 = 0.116$. These associations are illustrated in Supplemental Figure S4. In other words, neither the magnitude of switching costs nor their change across alertness states accounted for the corresponding variation in Miller's bound violation.

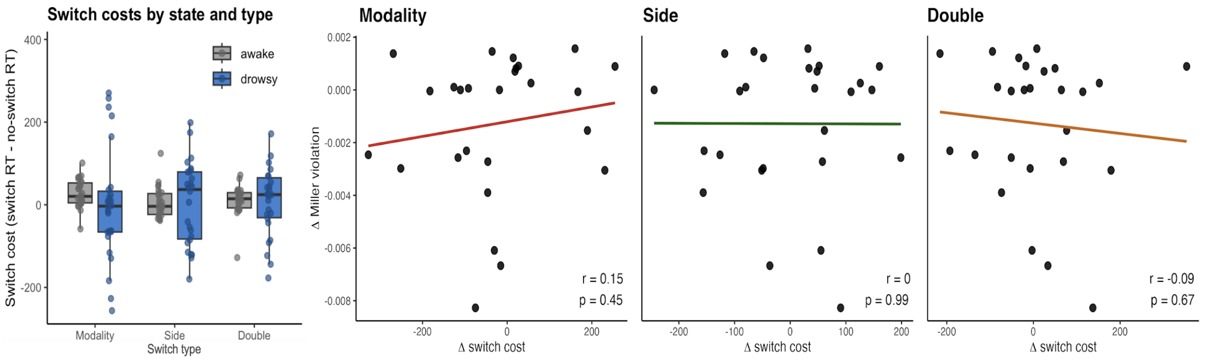


**Supplemental Figure S4**. Switch costs by alertness state and their relationship with changes in Miller's bound violation. Left: distribution of switch costs (relative to no-switch trials) for modality-only, side-only, and double-switch conditions in awake and drowsy states. Right: correlations between awake-to-drowsy changes in switch costs and changes in Miller's bound violation, showing no reliable associations.

Another potential concern when interpreting race model violations is that they may be influenced by overall response slowing or by changes in unisensory processing, rather than reflecting genuine multisensory interactions (Couth et al., 2018). This is particularly relevant here given the marked slowing observed during drowsiness.We first assessed whether global RT slowing across participants was associated with changes in Miller's bound violation. The awake-to-drowsy change in mean RT was not significantly correlated with the corresponding change in violation ($r = -9.37$, $p = 0.062$), indicating that overall slowing does not explain the observed pattern.

We then examined whether changes in unisensory RTs could account for variation in violation. Pearson correlations revealed that increases in auditory RTs were significantly associated with decreases in violation ($r = -0.41$, $p = 0.036$), while a similar but non-significant trend was observed for tactile RTs ($r = -0.34$, $p = 0.093$). The change in mean unisensory RT also showed a significant negative association ($r = -0.40$, $p = 0.044$), whereas the relationship with the fastest unisensory RT did not reach significance ($r = -0.32$, $p = 0.108$). These results were confirmed using Spearman correlations (auditory: $\rho = -0.43$, $p = 0.029$; mean unisensory RT: $\rho = -0.41$, $p = 0.035$), with no significant effects for tactile or fastest RT measures (all *p*s > 0.11). To further characterise this relationship, participants were divided based on whether they exhibited higher violation values in the drowsy state relative to wakefulness. Those showing reduced violation in drowsiness exhibited significantly greater slowing in auditory RTs ($t(19.84) = 2.88$, $p = 0.009$), as well as in mean unisensory RTs ($t(22.19) = 2.33$, $p = 0.030$) and fastest unisensory RTs ($t(23.46) = 2.15$, $p = 0.042$), with no significant difference observed for tactile RTs ($t(23.68) = 1.59$, $p = 0.125$). Finally, regression analyses confirmed that unisensory RT changes did not reliably explain variance in the change in violation when considered jointly ($F(2,23) = 2.40$, $p = 0.113$, $R^2 = 0.17$). A model including mean unisensory RT alone reached significance ($\beta = -7.68 \times 10^{-6}$, $p = 0.044$), but this effect was small and consistent with the negative associations reported above. A similar model based on the fastest unisensory RT did not reach significance ($F(1,24) = 2.80$, $p = 0.108$). Together, these results indicate that the observed pattern of race model violation cannot be attributed to global slowing or to changes in unisensory processing.

#### *ERP difference-in-differences analysis across alertness states*

The differences between the event-related potentials elicited by the multisensory stimulus and the linear sum of the corresponding unisensory responses were first estimated within each alertness state and are shown in Figure 2 (manuscript). We then tested whether the effects traditionally associated with multisensory integration, such as super- or subadditivity signatures, were modulated by alertness by directly comparing awake and drowsy muti-sum difference waveforms using a cluster-based permutation paired-samples *t*-test approach. This approach isolated spatiotemporal clusters in which multisensory vs. sum differences varied as a function of state, while controlling for multiple comparisons across electrodes and time.

Direct comparisons between awake and drowsy conditions (multi–sum ERP difference in differences) revealed that the subadditivity effect found in later stages of processing was significantly reduced during drowsiness in two electrodes clusters: positive (160–388 ms, $p = 0.003$) and negative (192–420 ms, $p = 0.003$), as represented in Supplemental Figure S5.

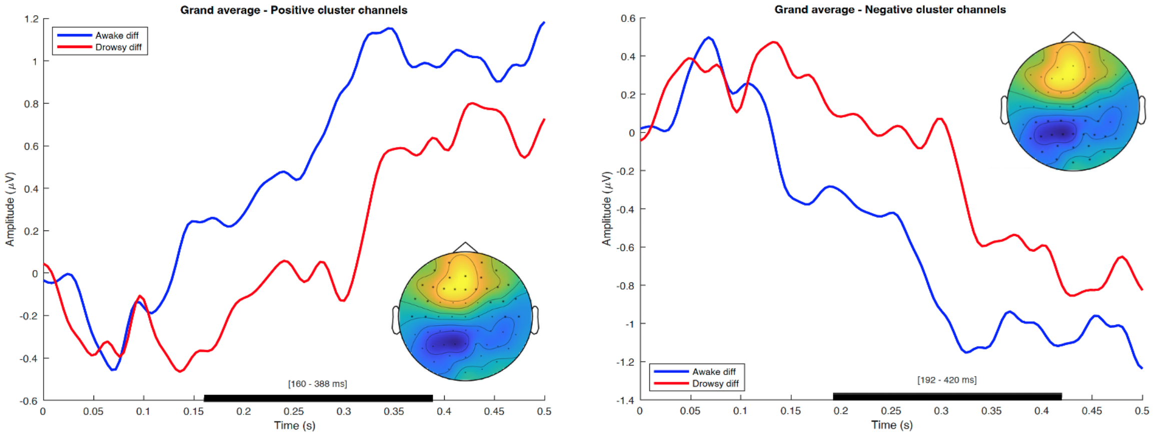


**Supplemental Figure S5**. Grand-average difference-waveforms (multisensory – summed unisensory responses) for awake (blue) and drowsy (red) trials, plotted over channels belonging to the positive (left) and negative (right) clusters identified in the cluster-based permutation analysis where differences between states were statistically significant. Shaded horizontal bars denote significant time windows. Topographic insets illustrate the spatial distribution of the respective clusters.

#### *Peak latency control analysis*

Cluster-based permutation tests identify time windows showing reliable condition differences but do not directly test whether ERP component peaks occur at different latencies. As suggested during peer-review, we performed an additional peak-latency control analysis on the multisensory interaction waveform (multisensory − sum of unisensory responses). For each participant, we averaged the ERP across the electrodes belonging to the early and late clusters and extracted the latency of the most prominent peak within the corresponding time window. In the early window (40–150 ms), we quantified the negative peak (fronto-crental cluster with lowest *p*-value); in the late window (180–450 ms), we quantified the positive peak (parieto-occipital cluster with lowest *p*-value). Peak latencies were then compared between wakefulness and drowsiness using paired-sample *t*-tests.

Paired-sample *t*-tests revealed no significant differences in peak latency between wakefulness and drowsiness in either time window. In the early cluster, peak latency did not differ between states, $t(25) = -1.16$, $p = 0.256$ (awake: 93.08 ms, SD = 34.16; drowsy: 104.62 ms, SD = 32.78). In the late cluster, peak latency again did not differ between states, $t(25) = -0.25$, $p = 0.805$ (awake: 251.08 ms, SD = 64.83; drowsy: 255.69 ms, SD = 65.94). These findings suggest that the cluster-based results are better explained by differences in the magnitude and statistical emergence of multisensory effects, rather than by systematic shifts in ERP peak latency.
